# Targeting NEDD9-SH3 with a Covalent Peptide Controls Endothelial Phenotype

**DOI:** 10.1101/2025.07.10.663547

**Authors:** Andriy O. Samokhin, Hyuk-Soo Seo, Alison Leed, Behnoush Hajian, Gregory H. Bird, David C. McKinney, Progyaparamita Saha, Jacqueline Daum, Jamie A. Moroco, Jenna Yehl, David Kornfilt, Anna Szczeniowski, Elyse Petrunak, Steven Horner, Jay H. Kalin, Nancy Leymarie, Virendar K. Kaushik, Loren D. Walensky, Philip A. Cole, William M. Oldham, Matthew L. Steinhauser, Sirano Dhe-Paganon, Bradley A. Maron

## Abstract

Src homology 3 (SH3) proteins regulate numerous fibroproliferative pathophenotypes including pulmonary arterial hypertension (PAH) but are challenging to target therapeutically. We innovated a peptidomimetic that occupies the canonical focal adhesion kinase (FAK) binding site on the SH3 domain of the neural precursor cell expressed, developmentally down-regulated 9 (NEDD9) protein, a pro-PAH regulator. Peptidomimetic derivatization with a bromoacetamide group alkylated a NEDD9 cysteine positioned uniquely among SH3 domains (Cys18), which stabilized the RT loop, prevented FAK binding, and inhibited human pulmonary artery endothelial cell (HPAEC) migration. When linked to a thalidomide moiety, the peptide showed degrader activity of NEDD9 protein and, therefore, we next investigated therapeutic application of NEDD9 inhibition. In HPAECs, si-NEDD9 downregulated sulfatase-1, which increased podosome rosette formation and cell migration via 6-O-desulfation of glycocalyx-forming heparan sulfate proteoglycans, and reversed vascular remodeling and PAH *in vivo*. Whereas sulfatase-1 overexpression decreased pulmonary endothelial podosome formation, cell migration, and tube formation and increased collagen III synthesis, sulfatase-1 knockdown prevented fibroproliferative remodeling and pulmonary hypertension in PAH *in vivo*. These data leverage cysteinyl thiol reactivity to establish an SH3 domain-targeting structure-validated covalent peptide and identify two convergent mechanisms through NEDD9 that control endothelial phenotype, including reverse remodeling via sulfatase-1 transcriptional control. Overall, this study advances an SH3-specific therapeutic approach with relevance to PAH and other fibroproliferative pathophenotypes.

## Introduction

The Src homology 3 (SH3) domain proteins are highly conserved in eukaryotes and mediate numerous fundamental signaling events that control cellular homeostasis and intercellular communication.^1^ The SH3 proline-rich modular protein-binding domain can enable interaction with other proteins to form stable complexes, and events altering SH3-target interaction are implicated in the pathobiology of proliferative diseases.^2^ Structure-function analyses from experimental systems using mutant constructs imply that SH3 is a modifiable driver of abnormal cellular propulsion.^3,4^ However, canonical binding sites typical of SH3 lack pockets with exposed functionality and are mostly shallow, featureless surfaces.^5^ This hinders the development of target-specific therapeutics that affect cellular phenotype by interfering with functionally essential protein-protein interactions.^6^

Neural precursor cell expressed, developmentally down-regulated 9 (NEDD9) is a docking protein that in cancer cells interacts directly with focal adhesion kinase (FAK) via its SH3 domain to modulate cell phenotype through focal adhesions.^3,7^ Unlike nearly all other SH3 family proteins, NEDD9-SH3 contains a unique cysteinyl thiol within the RT loop at position 18, which we have shown functions as a redox sensor that controls fibroproliferative signaling pathways in human pulmonary artery endothelial cells (HPAECs).^8^ Computational models using structural data derived from p130Cas (BCAR1), which shares a high degree of homology with NEDD9-SH3, suggest that the residue at position 18 may be accessible from the canonical FAK binding pocket.^9^ Thus, we postulate that Cys18 could function as a molecular handle for a novel FAK peptidomimetic to NEDD9 (NEDDtide) to significantly increase its binding affinity, disrupt FAK-NEDD9 interactions and, in doing so, modulate HPAEC phenotype via focal adhesions.

Pulmonary arterial hypertension (PAH) is characterized by fibroproliferative remodeling of distal pulmonary arterioles that promotes early heart failure.^10^ This complex PAH pathophenotype is ascribed, in part, to an imbalance between anti-proliferative^11^ and pro-proliferative signaling pathways,^12^ consistent with divergent focal adhesion functionalities reported across experimental systems *in cellulo*.^13^ Indeed, FAK overactivation is reported in experimental PAH^14^ and NEDD9 gene silencing prevents PAH *in vivo*.^8^ Recently, however, we observed by bulk RNA-Seq that NEDD9 inhibition also decreases pulmonary endothelial sulfatase-1 (Sulf1), which is a hydrolase (extracellular endosulfatase) that regulates endothelial glycocalyx remodeling.^15–18^ Experimental cleavage of heparan sulfate proteoglycan disaccharides decreases focal adhesion size and enhances endothelial migration,^19,20^ whereas 6-O-sulfate (6OS) de-sulfation of heparan sulfate chains at [Glc/IdoA(2S)-GlcNS(6S)] by Sulf1 activity alters the heparan sulfate binding affinity to PAH growth factors and morphogens, such as WNT/β-catenin and TGF-β/BMP.^21–23^ These observations suggest an unrecognized functional relationship may exist between NEDD9-Sulf1 and focal adhesions in PAH.

Here, we used crystallography, protein mass spectrometry (MS), funnel-based drug screening methods, and cellular and *in vivo* disease models to test the hypothesis that selectively targeting NEDD9 using a NEDDtide modified with an electrophilic residue or Sulf1 silencing leads to differential effects on HPAEC phenotype. Our findings advance therapeutic development for SH3 proteins with implications for PAH and other morbidities characterized by disruption of focal adhesions functionality.

## Results

### Chemical modification of NEDD9-Cys18 by an acrylamide electrophile

Post-translational oxidative modification of Cys18 prevents NEDD9 proteasomal degradation by inhibiting the formation of a protein-protein complex with Mothers against decapentaplegic homolog 3 (SMAD3), leading to NEDD9-dependent fibrosis in HPAECs.^8^ We noted from a previously published human proteome-wide analysis developed to detect cysteine ligandability that N-[3,5-Bis(trifluoromethyl)phenyl] acrylamide **1** (**Fig. S1a**) causes subtle displacement of iodoacetamide at NEDD9-Cys18.^24^ To determine if **1** could exert an important biological effect on NEDD9 and, thus, support a drug discovery program leveraging Cys18, HPAECs were treated with vehicle (V) control or **1** (50-200 µM) for 30-120 min. Compared to V-treated cells, **1** decreased NEDD9 protein quantity by immunoblot under basal conditions (**Fig. 1a**) and restored NEDD9-SMAD3 complex formation in cells treated with H_2_O_2_ (500 µM) for 20 min to induce oxidant stress (**Fig. 1b,c**). Fragment **1** also decreased fibrosis *in cellulo* as measured by anti-collagen III immunofluorescence (**Fig. 1d**).

**Figure 1.**
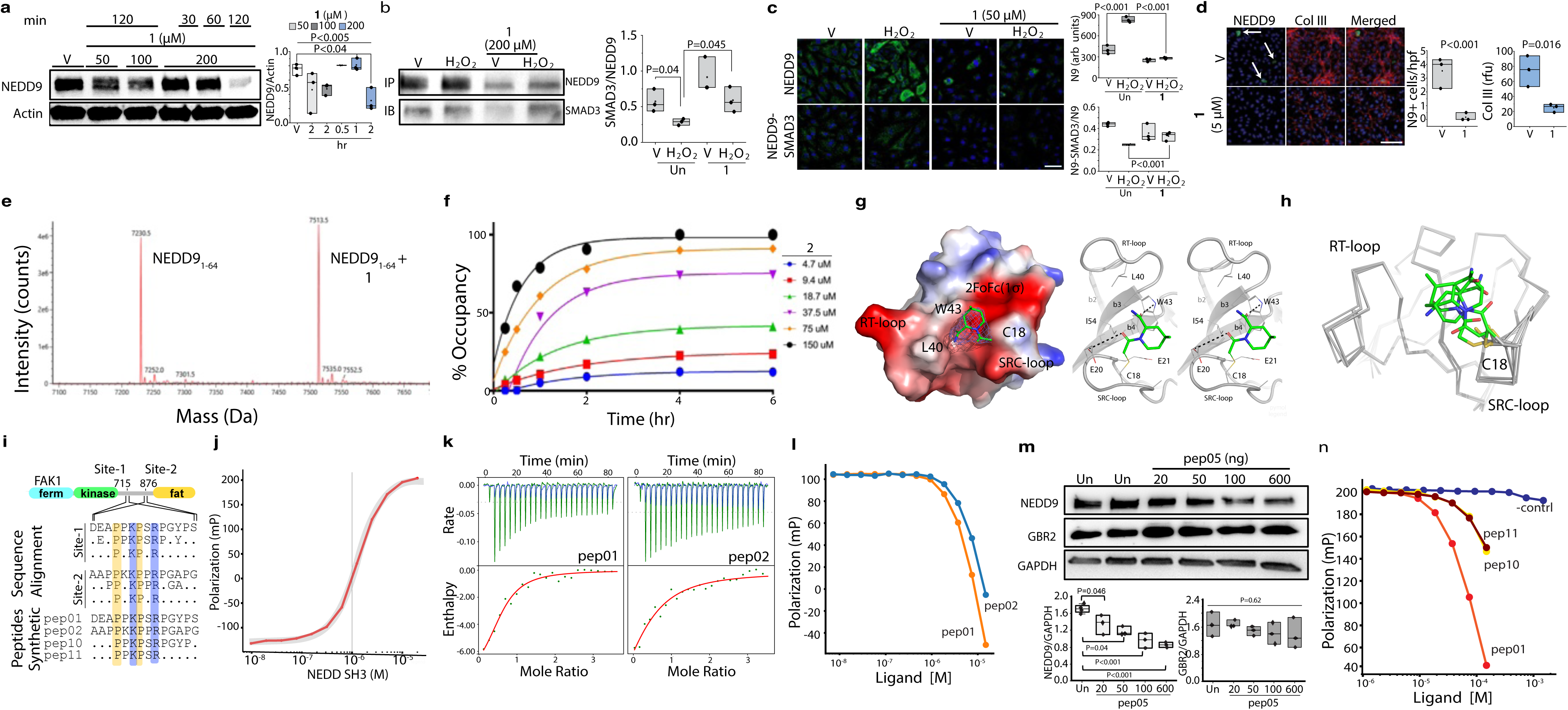
NEDD9-SH3 target engagement with a covalent electrophile and peptide. **a.** Dose- and time-course effect of Fragment **1** on NEDD9 expression in cultured human pulmonary artery endothelial cells (HPAECs) (*n*=3). **b.** Anti-NEDD9 immunoprecipitation (IP), anti-SMAD3 immunoblot (IB) was performed on HPAECs treated with H_2_O_2_ (500 μM) for 20 min in the presence or absence of **1** (*n*=3). **c.** The effect of **1** on the NEDD9-SMAD3 protein-protein interaction in H_2_O_2_-treated HPAECs measured by proximity ligase assay and quantitated relative to overall NEDD9 expression (*n*=3). Scale bar, 40 μm; N9, NEDD9. **d.** The effect of **1** for 24 hr on cellular NEDD9-positivity (N9+) and collagen III (Col III) deposition in HPAECs measured by immunofluorescence (*n*=3). Scale bar, 50 μm. **e.** Intact mass spectra demonstrating a modification of purified NEDD9-SH3 by **1** at Cys18. **f.** Dose-response and time-course effect of treatment with **2** on NEDD9-SH3 shows occupancy at Cys18 measured by MS2. **g.** [left] Electrostatic surface representation of the **2** – NEDD9-SH3 complex. The associated electron density is displayed at the 1σ level. The compound binding site is labeled. [right] Stereoscopic ribbon diagram of **2** in complex with NEDD9-SH3, highlighting the covalent bond between the acyl amide of **2** and Cys18. Hydrogen bonds between **2** and residues Trp43 and Glu20 are also shown. **h.** 3-D alignment of all 7 protein molecules in the asymmetric unit demonstrates three poses of **2** shown in stick format. **i.** Sequence alignment includes Site-1 and Site-2 of human FAK1 centered on Pro715 and Pro876, respectively, highlighting the consensus motif (PxKPxR). All synthetic peptides are N- and C-terminally capped. **j.** Fluorescence polarization measurements showing binding of FITC-labeled NEDDtide probe (FITCylated pep01) to GST-fused construct of the NEDD9-SH3 domain. The sigmoidal curve reflects the fitted binding model, indicating a dissociation constant (K_D_) of 4 µM. **k.** Isothermal titration calorimetry (ITC) experiments were conducted to determine the binding affinity of two peptides (pep01 and pep02) to NEDD9. (Top) Raw ITC thermograms display heat release over time upon titration of each peptide into the NEDD9 solution. (Bottom) Integrated binding isotherms with the fitted binding model (red curve) used to determine the dissociation constant (K_D_). The binding affinity for pep01 was determined to be approximately 4 µM, whereas pep02 exhibited a weaker interaction with a K_D_ of approximately 14 µM. These results suggest that pep01 has a higher binding affinity to NEDD9 compared to pep02, supporting its potential as a more effective inhibitor. **l.** Determination of the IC50 of the NEDDtides pep01 and pep02 for disrupting the interaction between FITC-labeled pep01 and GST-NEDD9. **m.** The effect of HPAEC transfection for 10 min with NEDDtide pep05 on NEDD9 and GBR2, which is an SH3 protein that lacking the PxKPxR consensus motif and therefore served as a negative control (*n*=3-4/condition). **n.** Fluorescence polarization profiles the competitive displacement of FITC-labeled pep01 (50 nM) from NEDDtides pep01, pep10, and pep11 (sequences detailed in panel **i**).

A truncated form of NEDD9 that included only the SH3 domain (1-64) was analyzed by intact protein mass spectrometry, which confirmed a single modification by **1** (**Fig. 1e**) marked by the addition of 283 Da to the protein mass. Analysis of the SH3 domain protein with the alkylating agent, iodoacetamide, showed that only Cys18, and not Cys47 (the only other NEDD9-SH3 cysteine residue), is available for modification under the same conditions **(Fig. S1b)**. Similar findings were observed by 2-dimensional nuclear magnetic resonance (2-D NMR) (**Fig. S1c)**. To discover novel chemical matter that modifies NEDD9 Cys18, we utilized the SH3 domain-only construct as well as full length NEDD9 in a screening funnel that used mass spectrometry, x-ray crystallography and 2-D NMR (**Fig. S2**). Initial hits were found by screening N=6,120 covalent electrophiles against the NEDD9-SH3 domain by RapidFire-MS. Equimolar amounts of NEDD9-SH3 and the BRAF-Ras binding domain (RBD) were incubated with the fragment library as a means of performing the screen and counter screen simultaneously, as BRAF-RBD is an isolated domain with similar properties to the NEDD9-SH3 domain, including one reactive surface cysteine as well as 2 additional cysteines buried within the folded protein. Fragments that were found to adduct at least two-fold more to the NEDD9-SH3 domain were progressed to NEDD9 1-64 kinetic studies to further characterize the molecules. A dose- and time-dependent increase in adduct formation with (2*R*,4*R*)-1-(2-chloroacetyl)-4-methylpiperridine-2-carboxamide **2** (**Fig. S3a**) to NEDD9-SH3 was confirmed by intact MS (**Fig. 1f**, **Fig. S3b)** and 2-D NMR (**Fig. S3c)**, whereas deconvoluted MS did not show BRAF-RBD modification (**Fig. S3d**).

The structural basis by which **2** binds to NEDD9-SH3 was determined by co-crystallizing NEDD-SH3 domain with the compound. A 2.2 Å structure was resolved containing seven molecules in the asymmetric unit out of which six showed density for the compound. While the electron density for the covalent bond and the core of the piperidine ring were well-defined, the precise rotational orientation remained unclear (**Fig. 1g,h**). Nevertheless, in one rotational orientation, **2** approached Leu40 and Trp43, forming two hydrogen bonds within the domain pocket. One hydrogen bond occurred between the carbonyl of the amide group in the ortho position and the indole nitrogen of Trp43, while the other was formed between the acetamide oxygen and the side chain of Glu20. When analyzing full length NEDD9, we again observed a time-dependent increase in Cys18 alkylation by **2**; however, while this was the primary site of alkylation, as expected with this reactive electrophile, alkylation of Cys281 was observed at later time points and higher compound concentrations (**Fig. S4)**.

### Developing a NEDD9 chemical probe to optimize target engagement

To address the dilemma of SH3 domain target-selective drug delivery, we turned to substrate/binding-partner-based peptide chemical probes, and noted a yeast-display-derived consensus motif that is predicted to bind to NEDD9 with the sequence: PxKPxR.^25^ Proteome sequence analysis of established members of the NEDD9 signalome and interactome using Motif2^26^ revealed only one human gene (KIAA1522) with N*≥*3 occurrences of the motif and 10 human genes with exactly N=2 occurrences including PTK2 (FAK1) and PTK2B (FAK2), (**Fig. S5a**) (see methods for details). The motif occurs in the disordered linker between the kinase and four-helix-bundle domains of FAK1 and 2 (**Fig. 1i**).

Two 14 residue-long NEDDtides were synthesized (pep01 and pep02) corresponding to sites 1 and 2 of FAK1, respectively (see **Table S1** for a catalogue of all NEDDtides). Fluorescence polarization confirmed binding of the FITC-labeled pep01 probe to the GST-fused construct of the NEDD9 SH3 domain (**Fig. 1j**). The affinity of pep01 and pep02 to the SH3 domain of NEDD9 was demonstrated by isothermal titration calorimetry (ITC) that yielded K_D_ values of ∼4 μM and ∼14 μM, respectively, with high enthalpy values consistent with protein-protein interactions (**Fig. 1k**). Fluorescence polarization determined that the IC50 of NEDDtides pep01 and pep02 for disrupting the interaction between FITC-labeled pep01 and GST-NEDD9 was ∼1 μM (**Fig. 1l**).

To confirm target-engagement in cells, we developed pep03, which is a biotinylated version of pep01. Anti-NEDD9 and anti-biotin double immunofluorescence performed in HPAECs transfected with a scrambled (negative) control peptide (pep04) or pep03 demonstrated NEDD9-biotin colocalization at structures resembling focal adhesions (**Fig. S5b**). To develop a functional correlate to this observation, we next modified a shorter version of pep01 to a proteolysis-targeting chimera (PROTAC). The pep01 isoform was conjugated with a thalidomide moiety (pep05) (**Fig. S5c**) to engage E3 ubiquitin ligase-mediated proteasomal degradation.^27^ Compared to untreated HPAEC monolayers, transfection with pep05 for 10 min induced dose-dependent degradation of NEDD9 that was not observed for Growth Factor Receptor-Bound Protein 2 (GBR2) which is an SH3 domain protein lacking the PxKPxR consensus motif (**Fig. 1m**). We used these data as an indicator of successful target engagement and focused next on opportunities to improve the binding affinity of the NEDDtide, noting that comparatively shorter NEDDtides of 10 and 6 amino acids (pep10 and pep11, respectively) (**Fig. 1i**) retained most of the binding affinity of pep01 (**Fig. 1n**). We surmised that the synthesis of peptides containing electrophilic fragments would be able improve on this affinity by alkylating Cys18.

### Structure-guided chemistry leveraging Cys^18^ to develop an SH3 target-specific peptide

The FAK1 sequence (pep10) was docked in NEDD9-SH3 *in silico* using Alpha-Fold2^28^ and the highest ranked pose is shown in **Fig. 2a**. Our computational models of the FAK-NEDD9 interaction showed that the P_1_ and K_3_ residues straddled Arg59, and P_4_ and R_6_ straddled Trp43. Importantly, the co-crystal structure of Pep10 and BCAR1 (which shares 92% homology with NEDD9-SH3) indicated that K_3_ and R_6_ pointed in the same direction towards the RT-loop, forming hydrogen and ionic bonds with Glu17, Glu21, and Cys18 (**Fig. 2b,c, Fig. S6**).

**Figure 2.**
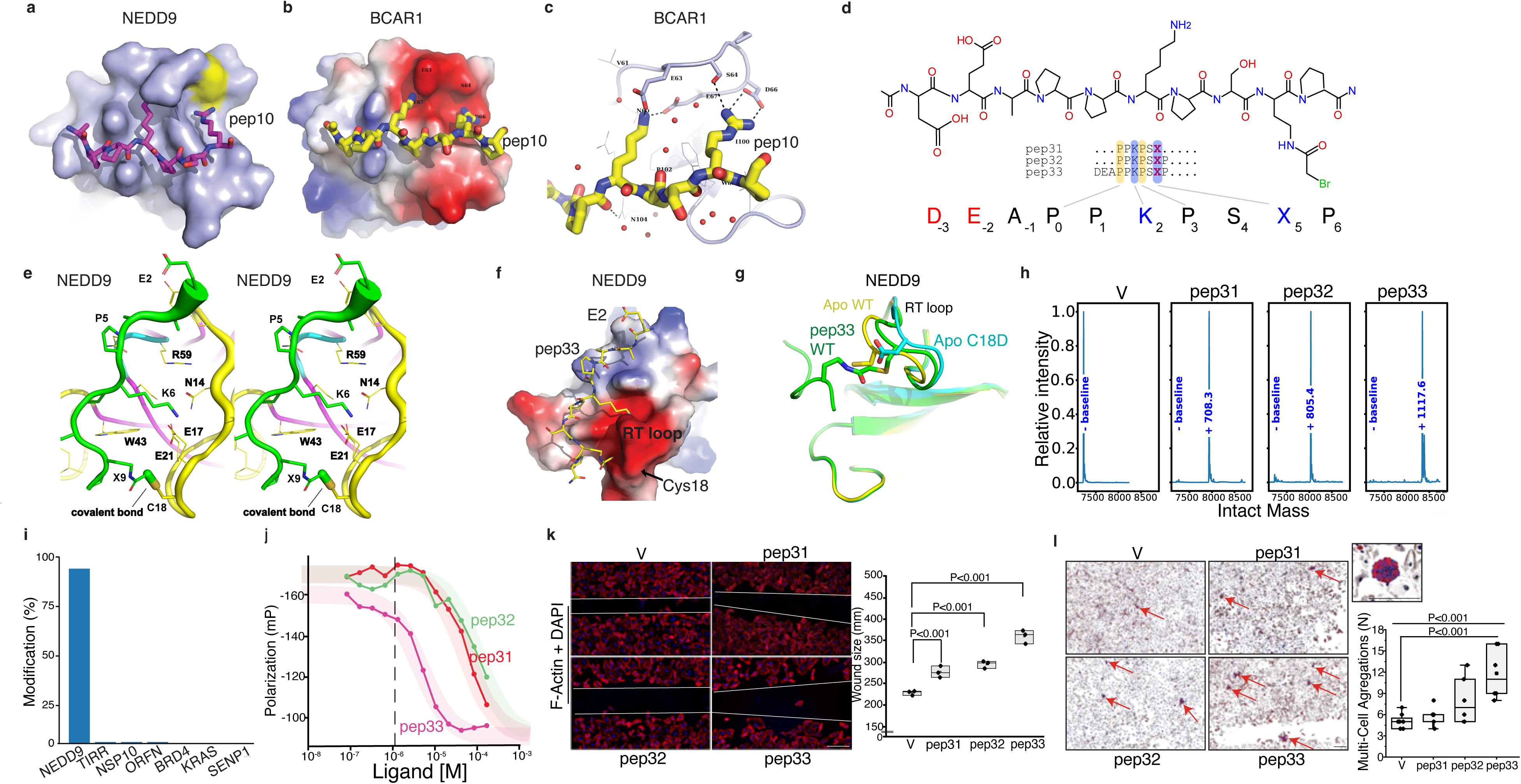
Leveraging NEDD9-Cys18 to develop a SH3 domain target-specific covalent peptide. **a.** The pep10 was docked *in silico* to produce multiple poses, the highest ranked of which is shown with a surface representation of the NEDD9-SH3 domain in light-blue and Cys18 in yellow. **b.** The structure of the SH3 domain of human BCAR1 (92% homology with NEDD9-SH3) was determined in complex with pep10, which contains the first PxKPxR motif of human FAK1. This electrostatic surface representation shows the complementarity of the charges between the positively charged basic peptide ligand and the negatively charged acidic BCAR1 binding pocket. **c.** Atomic closeup of the interaction between charged side chains (yellow, carbon atoms) and the SH3 domain showing ionic bonds and revealing the proximity of Arg to Ser64. Positively charged regions of BCAR1 are colored in blue, negatively charged regions in red, and neutral regions in white (−70 to +70 kT/e gradient). **d.** Sequence alignment of the covalent peptides pep31, pep32, and pep33 where X represents a bromoacetamide warhead. The chemical structure of pep33 is shown with positional labels below. **e.** Stereoscopic representation of the complex structure of NEDD9 with covalent pep33 in a cartoon format. Peptide side chains are shown as sticks, and nearby residues in NEDD9 are represented as lines. **f.** Surface representation of pep33 in complex with the SH3 domain of NEDD9. **g.** Ribbon overlay of the NEDD9-SH3 RT loop in the Apo WT structure (yellow), in the Apo D12A mutant that mimics oxidative modification at position 18 C18D (cyan), and in the pep33 NEDDtide complex structure (green). Pep33 is shown in ribbon format and Cys18 and Asp18 are shown in stick format. **h.** Deconvoluted intact mass spectra of the protein are shown after treatment with DMSO (vehicle control), pep31, pep32, and pep33. The DMSO-treated sample shows a single peak corresponding to the unmodified (baseline) protein mass. In contrast, samples treated with pep31, pep32, and pep33 exhibit additional peaks corresponding to mass shifts of +708.3 Da, +805.4 Da, and +1117.6 Da, respectively, indicating covalent adduct formation between the protein and each peptide. These mass increases are consistent with the expected mass additions for each peptide, suggesting specific and efficient covalent modification. Relative intensity is normalized across all conditions. The mass addition induced by NEDDtide modification of SH3 is provided adjacent to the spectra in blue. **i.** NEDDtide pep31 was analyzed using LC-MS against control proteins showing exceptional specificity for NEDD9. **j.** Determination of the IC50 of the NEDDtides pep31, pep32, and pep33 for disrupting the interaction between FITC-labeled probe and GST-NEDD9. **k.** Compared to vehicle (V) control, transfecting human pulmonary artery endothelial cells with NEDDtides pep31, pep32, and pep33 decreased cell migration proportionally to peptide length, assessed by wound healing assay (*n*=3). Blue, dapi; Scale bar, 400 μm. **l**. The effect of NEDDtides pep31, pep32, and pep33 on multi-cell cluster formation (arrows) (*n*=6-8). Box, representative cluster at high magnification. Scale bar, 600 μm.

To develop covalent NEDDtides, we noted the proximity of Arg6 to Cys18 and considered replacing the guanidinyl side chain of Arg6 with an electrophile. Given that MS screening and preliminary medicinal chemistry demonstrated that haloacetamides are stronger electrophiles to Cys18 compared to acrylamides, we opted to replace the guanidinyl group of Arg6 with a bromoacetamide. While aromatic or saturated rings linking the peptide backbone to the new electrophilic side chain could have enhanced reactivity, we observed no obvious feature that would hinder (or enhance) access, and thus chose a readily available aliphatic linker—bromoacetyl diaminobutanoic acid—for simplicity. The resulting peptide series (pep31-33) differed in overall peptide length (**Fig. 2d**, **Fig. S7a**).

A high-resolution crystal structure of pep33 in complex with NEDD9 revealed excellent electron density for the entire peptide, including the ethyl amide and the covalent bond with Cys18 (**Fig. 2e,f**). Importantly, the peptide adopted a canonical conformation within the NEDD9 pocket, consistent with our BCAR1/peptide structure, with an RMSD of ∼0.2 Å between the ligands. The position of Cys18 in the RT loop was slightly displaced compared to the apo structure, and the backbone atoms of the unnatural covalent side chain were ∼1.5 Å closer to Cys18, likely to accommodate the covalent bond.^28^ Site-directed substitution of Cys18 with aspartic acid demonstrates RT loop rotation under these oxidomimetic conditions, whereas engagement of SH3 with pep33 stabilizes the RT loop akin to the confirmation observed in the apo structure (**Fig. 2g**).

Intact MS analysis demonstrated that incubation with a 10-fold excess of pep31, pep32, or pep33 led to complete labeling of the SH3 domain of full length NEDD9 after 1 hr incubation at room temperature (**Fig. 2h**). We then assessed pep31 specificity by intact-MS analysis using six randomly selected cysteine-containing proteins and observed high specificity for NEDD9 (**Fig. 2i**). Fluorescence polarization assays with FITC-labeled pep01 showed strong competition after incubation for 1 hr at room temperature, with pep33 outperforming pep31 and pep32 by approximately an order of magnitude (**Fig. 2j**). This enhanced performance was likely due to the additional N-terminal DEA residues, which provided extra stabilizing interactions. Transfection of HPAECs with pep41 (which is pep33 modified with a thalidomide rather than a bromoacetamide residue) (**Fig. S7b**) for 2 hr induced a biphasic effect on NEDD9 degradation including a (subtle) paradoxical decrease in degradation at higher concentrations consistent with the ‘hook-effect’ observed in degrader-based assays (**Fig. S7c**).

Since FAK-NEDD9 interaction regulates focal adhesion assembly and, thus, cell movement, the biofunctionality of the NEDDtide series was tested in a wound healing assay using HPAECs. Compared to V-treated HPAECs, transfection with pep31, pep32, and pep33, which increase incrementally in length, caused a proportional increase in wound size (**Fig. 2k**) as well as the number of multi-cell clusters, which showed decreased migration and suggests decreased surface attachment and increased cell-cell adhesion^29^ (**Fig. 2l**).

### NEDD9 inhibition reverses PAH in vivo

We showed NEDD9 degradation with pep05 and pep41, and therefore next aimed to characterize therapeutic application of NEDD9 inhibition in PAH. To test this hypothesis, we used the SU-5416-Hypoxia-Normoxia rodent experimental model of PAH and administered si-NEDD9 to the rats using a disease-reversal treatment protocol (**Fig. 3a**). Compared to untreated rats, fibroproliferative pulmonary arterial remodeling and severe pulmonary hypertension were evident in the PAH model at day 14 when si-NEDD9 treatment was initiated. Intratracheal administration of si-NEDD9 beginning at protocol day 14 and resulted in significantly decreased PAEC and pulmonary artery smooth muscle cell (PASMC) proliferation, increased the pulmonary arterial lumen:vessel ratio, and improved right ventricular systolic pressure (RVSP) as well as RV mass (**Fig. 3b-e**) on day 32 without affecting mean arterial pressure (**Fig. S8a**) compared to treatment with scrambled (negative) control (Scr) siRNA. Reverse remodeling of pulmonary arterioles by NEDD9 inhibition was supported further by histopathological analysis using trichrome staining, picrosirius red staining (fibrillar collagen) (Fig. 8b) and immunostaining for collagen III (**Fig. S9a**).

**Figure 3.**
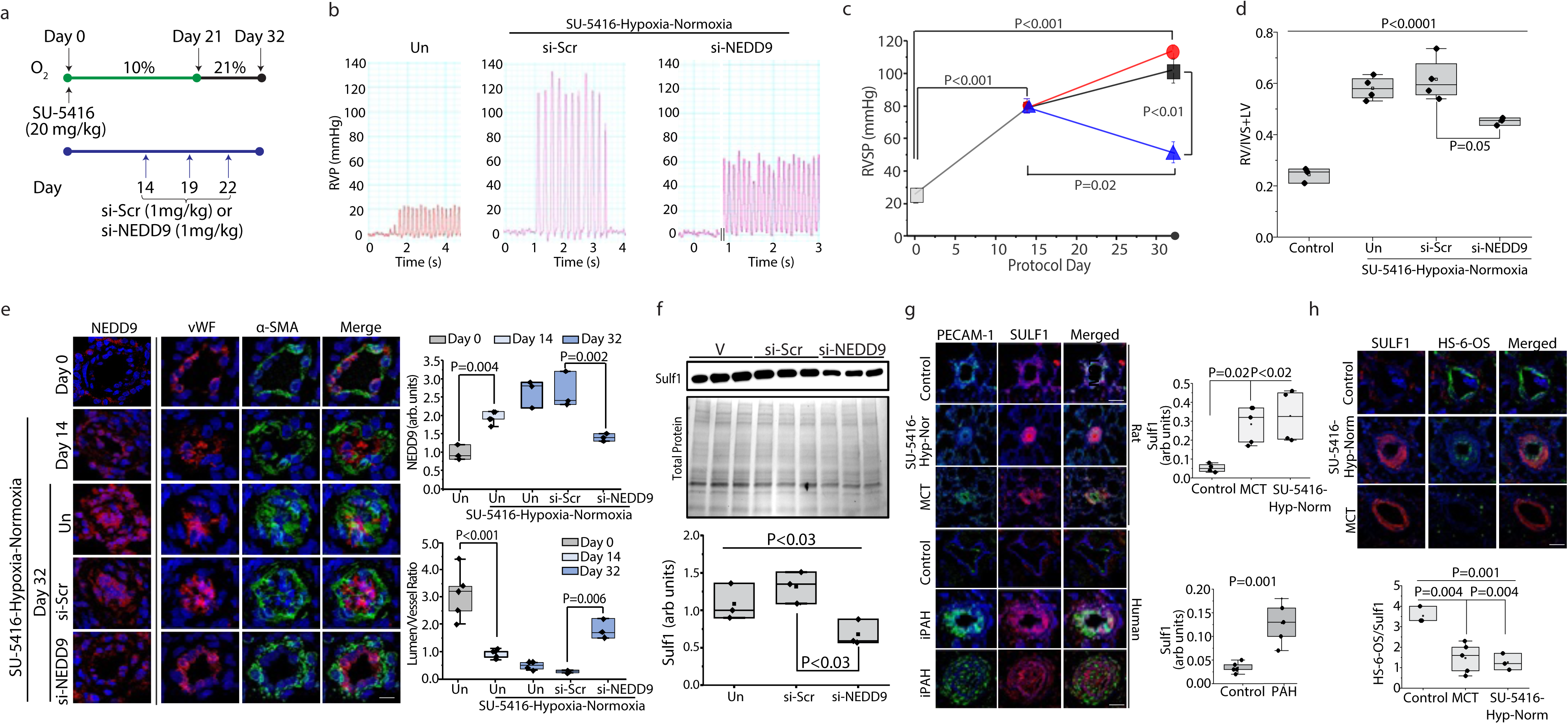
Transcriptional regulation of Sulf1 by NEDD9 is a reversible cause of PAH *in vivo*. **a.** Disease reversal protocol using the rat SU-5416-hypoxia-normoxia experimental model of PAH and timepoints for intratracheal administration of si-Scrambled (negative) control (Scr) or siRNA against NEDD9. **b.** Representative hemodynamic tracings for right ventricular pressure (RVSP) measured by invasive cardiac catheterization for each treatment condition. Un, untreated. **c.** The effect of si-NEDD9 on RVSP (*n*=3-5 rats/condition) and **d.** RV mass measured by Fulton index (*n*=3-4 rats/condition). Triangle, si-NEDD9 initiated at day 14; square, si-Scr initiated at day 15; circle, untreated. IVS, interventricular septum; LV, left ventricle. **e.** Paraffin-embedded lung sections isolated at key timepoints over the course of the PAH reversal protocol in each treatment condition were stained using anti-NEDD9, von Willebrand Factor (vWF), and *α*-smooth muscle actin (SMA) antibodies, and the immunofluorescence expression profile was analyzed for distal pulmonary arterioles (*n*=3-5 rats/condition). These data were used to quantify vascular NEDD9 expression and the lumen/vessel wall ratio (*n*=3-5 rats/condition). arb, arbitrary. Scale bar, 20 μm. **f.** Cultured human pulmonary artery endothelial cells were transfected with vehicle (V) control, si-scrambled (negative) control (Scr), or si-NEDD9 for 24 hr and Sulf1 expression was analyzed by immunoblot, standardized to total protein level (*n*=3). **g.** Paraffin-embedded lung sections from control, rat SU-5416-Hypoxia-Normoxia-PAH, and rat monocrotaline (MCT)-PAH (*n*=3-5 rats/condition) or isolated from idiopathic PAH (iPAH) patients (*n*=5/condition) at the time of lung explant were stained using anti-PECAM1 and anti-Sulf1 antibodies and the immunohistochemical profile in distal pulmonary arterioles was analyzed. Scale bar, 20 μm. **h.** Paraffin-embedded lung sections from control, rat SU-5416-Hypoxia-Normoxia-PAH, and rat monocrotaline (MCT)-PAH (*n*=3-5 rats/condition) were stained using anti-Sulf1 and anti-HS-6-OS antibodies and the immunofluorescence ratio in distal pulmonary arterioles was analyzed. Scale bar, 20 μm. HS, heparan sulfate.

### NEDD9 inhibition down-regulates Sulfatase-1

Impaired endothelial function following vascular injury has been shown to drive progressive fibroproliferative vascular changes in PAH, suggesting that reverse remodeling may require interventions which restore endothelial function.^30^ We re-interrogated pulmonary endothelial transcriptomic data from HPAECs that were untransfected or transfected with si-NEDD9 and treated with a PAH-relevant concentration of the pro-oxidant and pro-fibroproliferative hormone aldosterone.^8,31^ We observed that among top differentially regulated genes was Sulf1 (**Table S2, Fig. S9b**), and chose to study this further based on prior reports linking Sulf1 with growth factors that are involved in PAH and regulate FAK and cell migration.^21–23^ In cultured HPAECs, si-NEDD9 decreased protein expression (**Fig. 3f**) significantly compared to si-Scr, and reversed Sulf1 overexpression in remodeled pulmonary arterioles in SU-5416-Hypoxia-Normoxia-PAH *in vivo* (**Fig. S9b**).

Double immunofluorescence affirmed the relationship between NEDD9-Sulf1 and focal adhesions in HPAECs by showing co-localization of NEDD9-FAK (**Fig. S10a**) as well as Sulf1-NEDD9 (**Fig. S10b**), Sulf1-FAK (**Fig. S10c)**, and the Sulf1-paxillin, which is a focal adhesion marker (**Fig. S10d)**. We next used immunohistochemical analyses to demonstrate the relevance of these observations to endothelial dysfunction and noted that Sulf1 overexpression was evident in PECAM-1-positive cells *in situ* in both SU-5416-Hypoxia-Normoxia-PAH and the monocrotaline (MCT) (inflammatory) models of PAH, as well as lung sections explanted from patients with end-stage PAH (**Fig. 3g**).

### Heparan sulfate 6-O-desulfation in PAH

Post-translational modification of glycocalyx-forming heparan sulfate (HS) proteoglycans by Sulf1-dependent 6-O*-*desulfation is implicated in focal adhesion formation in cancer cells.^32^ Owing to phenotypic overlap between FAK in solid tumor cancer and fibroproliferative expansion of pulmonary arterioles in PAH,^14^ we next focused on the relationship between Sulf1 and HS-6-O*-*S in PAH. Compared to controls, immunofluorescence using an antibody recognizing the 6*-*O*-*S modification specifically^17^ indicated that upregulation of Sulf1 induced a reduction in the level of HS-6*-*O*-*S in SU-5416-Hypoxia-Normoxia-PAH and MCT-PAH *in situ* (**Fig. 3h**), and correlated inversely with vascular Sulf1 and NEDD9 (**Fig. S11a,b)**. Further, pulmonary endothelial levels of the transmembrane HS proteoglycan neuropilin-1 (NRP1), which functions as a co-receptor to pro-angiogenic VEGFR2 and stabilizes its interaction with VEGF in endothelial cells, was increased in PAH *in vivo* and in idiopathic PAH patients (**Fig. 4a**), suggesting 6*-*O-desulfation of HS attached to NRP1 by Sulf1. In HPAECs co-cultured with PASMCs directly, transmission electron microscopy showed colocalization of immunogold labeled anti-NRP1 and anti-HS-6*-*O*-*S antibodies, with the greatest signal at areas resembling focal adhesion (**Fig. 4b**).

**Figure 4.**
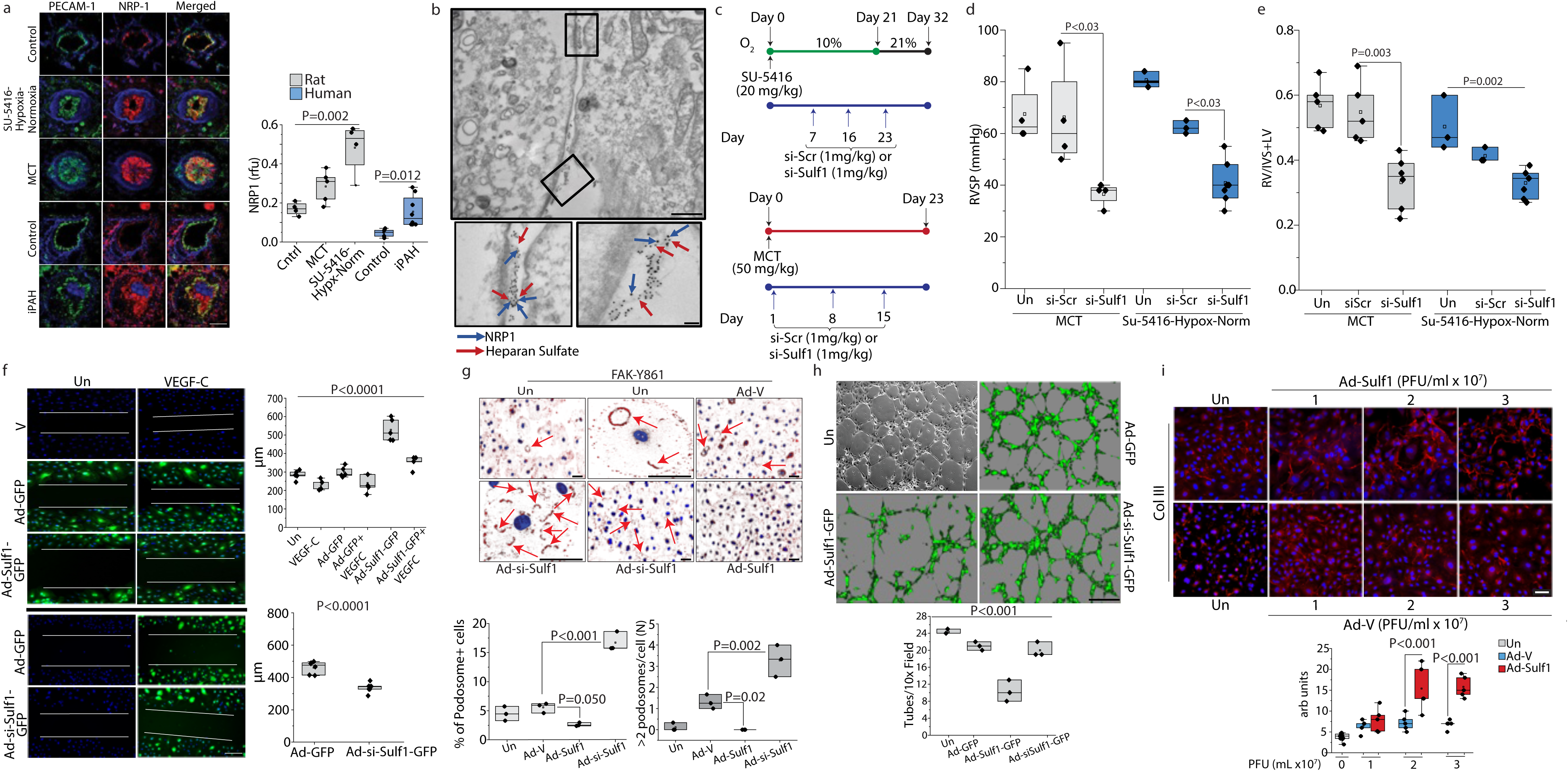
Inhibiting HS-6O-desulfation by Sulf1 increases endothelial podosome rosettes formation and prevents PAH *in vivo*. **a.** Paraffin-embedded lung sections from control, rat SU-5416-Hypoxia-Normoxia-PAH, and rat monocrotaline (MCT)-PAH (*n*=4-5 rats/condition) or isolated from idiopathic PAH patients (*n*=5-8/condition) at the time of lung explant were stained using anti-PECAM1 and anti-NRP-1 antibodies and the immunohistochemical profile in distal pulmonary arterioles was analyzed. Scale bar, 20 μm. **b.** Human pulmonary artery endothelial cells (HPAECs) and human pulmonary artery smooth muscle cells were co-cultured in a barrier-free system to recapitulate the direct cell-cell contact conditions in PAH lesions. Double immunogold-labeled staining was performed using anti-NRP-1 and anti-Heparan sulfate antibodies, and imaged using transmission electron microscopy. Scale bar, 500 nm. Inset scale bar, 100 nm. **c.** Disease prevention protocol using the rat SU-5416-hypoxia-normoxia and monocrotaline (MCT) experimental models of PAH and timepoints for intratracheal administration of si-Scrambled (negative) control (Scr) or siRNA against NEDD9. **d.** The right ventricular systolic pressure (RVSP) measured by cardiac catheterization (*n*=3-7 rats/condition) and **e.** RV mass measured by Fulton Index. IVs, interventricular septum; LV, left ventricle. **f.** Human pulmonary artery endothelial cells were untreated or treated with VEGF-C (100 nM) for 6 hr and transfected with vehicle (V) control or adenovirus (Ad) containing green fluorescent protein (GFP), Sulf1 cDNA, or siRNA-Sulf1, and cell migration was quantitated using the wound healing assay. Black line, separate experiment iteration without VEGF-C or other treatment (*n*=4-7). Blue, DAPI. Scale bar, 400 μm. **g.** HPAECs were untransfected or transfected with an Ad carrying vehicle (V) control, Scr, Sulf1, or si-Sulf1 and stained with an anti-FAK-Y861 antibody. The number of podosome rosette-positive cells (podosome+) per low power field (40x) and the number of cells with >2 podosomes was quantitated (*n*=3). Arrows indicate podosome structures. Scale bar, 50 μm. **h.** The number of tubes formed per 10x field was counted for untreated HPAECs and cells transfected with Ad carrying V, Sulf1, and si-Sulf1 (*n*=3). Scale bar, 300 μm. **i**. The effect of Ad-Sulf1 on collagen III deposition in HPAECs (*n*=5-10). PFE, plaque forming unit (N=1-3 PFEs). Scale bar, 50 μm.

### Sulf1 knock down improves fibroproliferative remodeling and PAH in two models in vivo

Disease-prevention treatment protocols were used to detect the effect of intratracheal siRNA-Sulf1 (1 mg/kg) or si-Scr (1 mg/kg) administration on pulmonary hypertension in the rat MCT-PAH and SU-5416-Hypoxia-Normoxia PAH experimental disease models (**Fig. 4c**). Inhibition of Sulf1 expression in remodeled pulmonary arterioles by si-Sulf1 caused a significant decrease in RVSP, RV mass, collagen deposition and fibroproliferative remodeling compared to treatment with si-Scr (**Fig. 4d,e, Fig. S12**).

### Sulf1 regulates HPAEC migration

Using a wound healing assay, we found that PAH-HPAEC migration is decreased significantly compared to donor controls (**Fig. S13a**), which was directionally similar to the effect of VEGF inhibition on angioproliferation in PAH, as reported by others.^33^ Compared to untransfected cells, adenovirus (Ad)-mediated Sulf1 overexpression attenuated the effect of VEGF-C on HPAEC migration whereas si-Sulf1 increased cellular mobility (**Fig. 4f**).

FAK is pivotal for focal adhesion assembly and turnover, and FAK activity is regulated by phosphorylation at several tyrosine residues including Y576 located in the kinase domain and Y861 that has been proposed to regulate FAK-NEDD9 interaction.^34^ In HPAECs probed for FAK-Y576 and FAK-Y861 expression by immunofluorescence, we noted the formation of podosome rosettes, which are actin-based dynamic microdomains that contain FAK and regulate cell migration via specialized extracellular matrix-degrading enzymes.^35,36^ To confirm that these structures were indeed podosomes, we performed immunostaining for cortactin, vinculin, NRP1, and F-actin, and noted staining patterns akin to FAK-Y576 and -Y861 (**Fig. S13b**). Owing to our findings that Sulf1 co-localizes with NEDD9 and FAK in HPAECs (Fig. S10), we postulated that modulating Sulf1 expression could therefore affect podosome rosette formation. Compared to untransfected cells or Ad-Scr controls, Ad-si-Sulf1 and Ad-Sulf1 caused a significant increase and decrease, respectively, in the number of podosome rosettes in HPAECs (**Fig. 4g**), whereas Sulf1 inhibition also transformed the rosette phenotype from circular to crescent-shaped (Fig. S13b).

### Sulf1 overexpression regulates the angiogenic and fibrotic potential of HPAECs

Considering the effect of Sulf1 on HPAEC mobility and podosome rosette formation, we next examined the role of Sulf1 on angiogenesis. Compared to transfection with Ad-Scr, Ad-Sulf1 inhibited capillary tube formation (**Fig. 4h**) that was similar in magnitude to the difference in tube formation observed for PAH-HPAECs versus human control donor HPAECs (**Fig. S13c**). To determine if the anti-remodeling effects of Sulf1 inhibition on pulmonary arterioles *in vivo* involved changes in endothelial fibrillar collagen deposition *per se*, cultured HPAECs were also analyzed using anti-collagen III immunofluorescence, which showed that Ad-Sulf1-transfected HPAECs had increased collagen III deposition compared to control (**Fig. 4i**).

## Discussion

Structure-guided analyses were used to develop a bromoacetamide-modified NEDDtide series that reacts with Cys18, expressed at a position on the flexible RT loop unique to NEDD9, to address the long-standing challenge of SH3 domain-targeting for drug discovery. The NEDDtide displaced a FAK probe *in vitro* to inhibit the focal adhesion-mediated process of endothelial migration *in cellulo*, whereas degrader activity could be induced by NEDDtide modification with thalidomide. The therapeutic potential of NEDD9 inhibition was supported by experiments demonstrating that si-NEDD9 decreased Sulf1 expression, which regulates pulmonary endothelial 6-O-desulfation of glycocalyx-forming heparan sulfate proteoglycans. Inhibition of Sulf1, in turn, increased podosomes and induced migration in HPAECs, suggesting that reverse remodeling observed by si-NEDD9 in PAH *in vivo* was determined, in part, by normalizing the endothelial phenotype via NEDD9-Sulf1 signaling. Taken together, these findings advance a novel SH3 domain-targeting technology and show differing effects by pharmacological or transcriptional approaches that target focal adhesion components through NEDD9, with direct relevance to PAH drug-development programs.

Consistent with other SH3 domains, the NEDD9-SH3 apo structure includes flat surfaces that typically hinders small molecule target-specific drug discovery.^37^ To address this challenge we used a peptide approach, which provides high target specificity but is also associated with low potency and low cell-permeability. A bioinformatics analysis identified a conserved proline rich motif (PxKPxR) at the FAK binding cleft of NEDD9-SH3. Our *in vitro* assays indeed confirmed that a series of NEDDtides interacted with NEDD9-SH3 at favorable K_D_; however, we reasoned that limited potency could ultimately hinder progressing the NEDDtides into cellular (and ultimately *in vivo*) assays. Thus, we focused on developing an electrophilic warhead that would take advantage of Cys18, an established redox sensitive thiol^8^ and essentially unique to NEDD9 (i.e., of 221 SH3 proteins, CASS4 is the only other SH3 domain-containing protein expressing cysteine at position 18).^38^

From the complex crystal structure involving the unmodified NEDDtide and BCAR1, which shares substantial homology with NEDD9 and is also implicated in PAH pathobiology,^39^ we anticipated that the terminal arginine residue within the binding motif was near Cys18, and used this feature to direct further chemical modifications with a warhead. This approach led to a unique covalent peptide that leveraged the binding strength of a covalent bond with high specificity of non-covalent interactions enabled by the amino acid sequence of the consensus motif, limiting off-target effects. As screening efforts had identified that a small molecule with a chloroacetamide warhead could form a specific covalent bond with Cys18, we modeled various NEDDtide modifications, ultimately settling on a bromoacetamide modified di-aminobutyric acid residue as a warhead to establish a covalent bond between NEDDtide and Cys18. Taken together, this establishes to the best of our knowledge the first-ever structure-validated covalent peptide ligand targeting an SH3 domain.

Oxidation of NEDD9-Cys18 has been shown to impair a critical protein-protein interaction with SMAD3, leading to pulmonary endothelial fibrosis.^8^ Here, site-directed substitution of cysteine with aspartic acid, functioning as a surrogate oxidomimetic,^40^ induced RT loop repositioning. Others have demonstrated that RT loop flexibility enables conformational changes in ordered protein regions that regulates protein function.^41^ We observed that RT loop repositioning under oxidomimetic conditions was subtle; however, it is notable that the bromoacetamide-modified NEDDtide stabilized the RT loop in an orientation akin to the apo structure, and covalent interaction between **1** and Cys18 preserved NEDD9-SMAD3 protein-protein interaction under oxidizing conditions *in cellulo*. These findings implicate functional consequences of NEDD9 RT loop rotation, regulated by redox sensing capability of Cys18, and that Cys18 stabilization with a covalent fragment is a suitable potential target for molecular glue-based technologies that aim to maintain NEDD9 homeostasis.

The NEDDtide series inhibited the interaction of a FAK probe with NEDD9-SH3, providing key empirical evidence validating prior models predicting that a conserved PxKPxR motif within the peptide recognition module^42^ could be leveraged to develop an asset that interacts with Class VII SH3 proteins.^25^ This effect appeared contingent upon the conserved binding motif, since degrader activity was not observed for GBR2, an SH3 protein lacking the consensus sequence.^25^ Target engagement and consequential biological relevance of these findings were confirmed by showing that cellular migration was inhibited directly by the bromoacetamide-modified NEDDtide. This observation aligns with data indicating that the FAK-NEDD9 protein-protein interaction is essential for focal adhesion assembly,^43^ which underlies cellular propulsion in proliferative disease states including PAH^44^ and solid tumor metastasis.^45^ Thus, taken together, our findings illustrate specific therapeutic potential achieved by interventional approaches affecting focal adhesions through NEDD9, the target of FAK.

Modification of the NEDDtide with a thalidomide residue decreased NEDD9 protein quantity, likely through E3 ligase engagement as reported for other similar PROTAC technologies.^46^ Indeed, therapeutically-modifying NEDD9 bioavailability has important implications for diseases characterized by dysregulated cellular proliferation, as dominant negative NEDD9 genotype has been shown previously to resist osteoclastogenesis,^47^ metastatic mammary cancer,^48^ and rheumatoid arthritis,^49^ for example, and in this study, si-NEDD9 reversed hypertrophic and fibrotic remodeling of distal pulmonary arterioles in PAH *in vivo*. The differential effect on cell migration we observed by disrupting FAK-NEDD9 protein-protein interactions with the NEDDtide *versus* NEDD9 gene silencing reinforces a framework in which the PAH phenotype is ultimately determined by imbalance between occlusive and longitudinal vascular proliferation, representing pathologic and adaptive injury response patterns, respectively.^50^ Niihori and colleagues^51^ and others^52^ have reported on the importance of impaired angiogenesis following vascular injury as a fundamental pathogenetic mechanism accounting for the obliterative vasculopathy of PAH. Our data reinforce and extend this framework by demonstrating that dual regulation of these processes can occur by affecting a single intermediary, NEDD9 itself, through genomic or pharmacologic interventions.

It has long been recognized that heparan sulfate proteoglycans (HSPG) regulate extracellular matrix structure and other fundamental endophenotypes underlying pulmonary hypertension, and recent data suggest vascular compartment-specific HSPG patterns that differ across disease subgroups.^53^ However, until now little has been reported on the pathogenetic role of HSPG in PAH *per se*. Our efforts to study the salutary effects of si-NEDD9 on the PAH pathophenotype suggested inhibitory transcriptional regulation of Sulf1 by NEDD9. Upregulation of Sulf1, in turn, has been implicated in proliferative pathophenoptypes including various adenocarcinomas, but beforehand was linked to PAH only indirectly through disease-relevant molecular targets including Wnt, BMP, and VEGF.^21–23^ Here, Sulf1-dependent 6-O-desulfation of HSPG diminished VEGF-dependent endothelial migration, decreased endothelial tube formation, and increased fibrosis. Thus, another important finding from this work is that programmed and target-specific desulfation is a novel and modifiable molecular mechanism underlying endothelial dysfunction that controls adverse vascular remodeling in PAH.

Numerous proteoglycans have been reported as substrates in extracellular matrix remodeling, including versican as well as agrin that is overexpressed in PAH pulmonary endothelial cells.^54^ We noted that NRP1 is a cell surface proteoglycan with several known functions linked to PAH pathobiology including cell migration and angiogenesis,^55^ and endothelial cells preferentially express NRP1 with heparan sulfate attached.^56^ It has been shown that HS attachment to NRP1 regulates co-receptor function for VEGF-VEGFR2 interactions as well as for the interactions of NRP1 with integrins and focal adhesion proteins.^57^ Those data suggest that HS bound to NRP1 is a substrate for Sulf1, which is supported empirically by our findings using dual immunolabeling transmission electron microscopy. It is notable also that this observation was made in a mixed HPAEC-HPASMC culture system lacking a mechanical or biological barrier, which is likely representative of internal elastic lamina-lacking PAH lesions, providing a basis to account for complex cell signaling patterns between heterologous and pro-migratory cell types observed at explanted PAH lungs *in situ*.

Analyzing the effect of Sulf1 bioactivity on the distribution of active FAK isoforms (phosphorylated at Y576 and Y861) revealed the presence of podosome rosettes, which are specialized actin-based dynamic microdomains of the plasma membrane that express a high concentration of matrix-degrading enzymes.^58,59^ Podosome rosettes share functions with focal adhesions and express overlapping molecular machinery, such as the F-actin ring, Vinculin, and FAK (but also express cortactin). Analyzing the expression of Sulf1 and NRP1 in HPAECs, we were able to detect similar circular structures but in significantly lower numbers compared to P-FAK and cortactin, whereas Sulf1 overexpression and knockdown with siRNA significantly decreased and increased cellular podosome rosette quantity, respectively. These differences suggested that Sulf1 overexpression attenuates podosome-dependent cell migration, which is pathological in PAH, but that normal migration can be restored through Sulf1 inhibition.

Alternative NEDDtide sequences were predicted to interact with the FAK binding grove of NEDD9 but were not tested empirically. Although NEDDtide-mediated interference of FAK-NEDD9 binding was internally consistent across several different assays *in vitro*, further data are needed to clarify the consequences of impaired FAK-NEDD9 interaction on FAK bioactivity in key cell types beyond HPAECs that promote vascular remodeling and known to express focal adhesions as well as in PAH *in vivo*. Similarly, experiments in this study were focused on establishing PROTAC target engagement and further biophysical profiling is needed to optimize its functionality. Recent data have suggested anabolic synthesis of macromolecular substrate fundamentally drives fibroproliferative remodeling in PAH.^60^ Taken together with wider availability of nanoscale imaging methods, this framework opens opportunity to explore atomic-level mechanisms underlying disease-reversal in PAH.^61^ MS-based quantitative profiling of HS-6-O-desulfation kinetics should be explored further to understand the specific implications of this enzymatic process to the disease state.

In conclusion, these data support the first-reported structure-enabled small molecule binder targeting an SH3 domain. A NEDDtide chemical probe is shown that utilizes the redox sensitivity of Cys18, demonstrating opportunity to leverage this technology as a degrader and protein-protein interaction disrupter for potential therapeutic purposes. We describe inhibition of the protein-protein interaction between FAK-NEDD9 and the linkage of NEDD9-Sulf1 signaling through 6-O-desulfation of HSPG as two convergent molecular mechanisms by which to modulate the pulmonary endothelial phenotype. We also demonstrate that this approach has direct implications on prevention and reversal of adverse pulmonary arterial remodeling and pulmonary hypertension *in vivo*. Overall, these data advance target-based therapeutics that affect pathogenic function of a crucial SH3 domain, with implications for PAH and other highly morbid proliferative pathophenotypes.

## Methods

### Synthesis of Fragment 1

*N*-(3,5-Bis[trifluoromethyl]phenyl) acrylamide.^1^ 3,5-Bis(trifluoromethyl)aniline (1.146 g, 5.0 mmol) was dissolved in anhydrous methylene chloride under argon (25 mL, 0.2 M) and cooled to 0 °C in an ice bath. Then, *N,N*-Diisopropylethylamine (0.969 g, 7.5 mmols) was added in one portion followed by the dropwise addition of acryloyl chloride (0.679 g, 7.5 mmols). The reaction was stirred for 2 h at 0 °C and monitored by thin layer chromatography (TLC). After completion, the reaction was quenched with water (1 mL), diluted in methylene chloride (20 mL), and washed with saturated sodium bicarbonate (2 x 100 mL). The organic layer was dried over anhydrous sodium sulfate, concentrated in vacuo, and purified by column chromatography (SiO_2_, 0-20% EtOAc/hexanes) to yield the desired product as a white, crystalline solid (0.922 g, 65%). ^1^H NMR (400 MHz, DMSO-*d*_6_): δ 10.78 (s, 1H), 8.31 (s, 2H), 7.75 (s, 1H), 6.40 (dd, *J* = 17.0 Hz, 9.5 Hz, 1H), 6.33 (dd, *J* = 17.0 Hz, 2.3 Hz, 1H), 5.86 (dd, *J* = 9.5 Hz, 2.2 Hz, 1H). ESI-LRMS calculated for C_11_H_7_F_6_NO: [M+H]^+^ = *m/z* 284.0, found: [M+H]^+^ = *m/z* 284.0 (**Fig. S1**).

Chemical reagents and solvents were purchased from Sigma-Aldrich and used as received. Reaction progress was monitored by TLC using pre-coated glass silica gel plates (SigmaAldrich F254, 60 Å, 250 µM). Column chromatography was carried out with a CombiFlash RF+ chromatography system (Teledyne ISCO) using pre-packed silica gel cartridges. NMR experiments were run on a Bruker 400 MHz (^1^H, 400 MHz; ^13^C, 100 MHz) spectrometer and processed using TopSpin 4.0 (Bruker). Chemical shifts (δ) are reported in parts per million (ppm) relative to the solvent residual peak and J-coupling constants (*J*) are reported in hertz (Hz). Multiplicity was designated with the following abbreviations: s (singlet), dd (doublet of doublets). Mass spectrometry was carried out using an Agilent SingleQuad LC/MSD instrument.

### Cell culture and treatments

Standard cell culture methods were used in this study, as reported previously by our laboratory.^2^ Human pulmonary artery endothelial cells and human pulmonary artery smooth muscle cells (all from Lonza) were grown to confluence using EBM-2 and SmGM-2 medium, respectively. All medium was supplemented with 5% fetal bovine serum (FBS); endothelial and SMC medium was also supplemented with a cell type-specific Bulletkit. All cells were incubated at 37°C, 5.0% CO_2_ and passaged twice-weekly using 0.5% trypsin/EDTA; experiments were performed on cell passages 4-10. The sex and demographics of cell types and commercially purchased reagents used in this study are provided in **Tables S3** and **S4**.

### PAH-HPAEC cell culture

Deidentified HPAECs isolated from idiopathic PAH patients (PAH-HPAECs) at the time of lung explant were provided graciously by the Pulmonary Hypertension Breakthrough Initiative, as reported previously.^2^ Experiments were performed on cell passages 4-10 (BWH IRB# 2015P001831).

### Treatment with Fragment 1

Fragment **1** was dissolved in DMSO (10 mM) which also served as the vehicle (V) control in chemical assays, and added to HPAEC cultures with final concentrations of 50, 100, and 200 mM for 30, 60 or 120 min before immunobloting.

### Proximity ligation assay

Duolink^®^ Proximity Ligation Assays (Sigma) was used according to the manufacturer’s instructions. For NEDD9 detection, proximity ligation was used to increase the signal compared to regular immunofluorescence and two different anti-NEDD9 antibodies raised in mouse and rabbit were used. To detect NEDD9-SMAD3 interaction, mouse anti-NEDD9 and rabbit anti-SMAD3 antibodies were used.

### HPAECs peroxide treatment

H_2_O_2_ (250 μM) was added to the cell culture medium at various time points and cells were incubated at 37°C, 5.0% CO_2_ until analysis.

### Cloning

All plasmids including mutants were prepared in pET21- or pET28-based vectors using standard cloning techniques with final expressed sequences as follows:

>NEDD9∼∼.0001-0064.KBBHOB.pHISSUMO MHHHHHHSDSEVNQEAKPEVKPEVKPETHINLKVSDGSSEIFFKIKKTTPLRRLMEAFAKRQGKEMDSLRFLYDGIRIQADQTPEDLDMEDNDIIEAHREQIGGMKYKNLMARALYDNVPECAEELAFRKGDILTVIEQNTGGLEGWWLCSLHGRQGIVPGNRVKLLI

>NEDD9∼∼.0006-0064.CRLJSA.pHH0103 MKIEEHHHHHHSSGKLMSPILGYWKIKGLVQPTRLLLEYLEEKYEEHLYERDEGDKWRNKKFELGLEFPNLPYYIDGDVKLTQSMAIIRYIADKHNMLGGCPKERAEISMLEGAVLDIRYGVSRIAYSKDFETLKVDFLSKLPEMLKMFEDRLCHKTYLNGDHVTHPDFMLYDALDVVLYMDPMCLDAFPKLVCFKKRIEAIPQIDKYLKSSKYIAWPLQGWQATFGGGDHPPKSDLEVLFQGPLSSGLVPRGSGTAAQPAMAQGALLMARALYDNVPECAEELAFRKGDILTVIEQNTGGLEGWWLCSLHGRQGIVPGNRVKLLIAAA

>BCAR1∼∼.0003-0071.PXAADH.pGEX6P2 MSPILGYWKIKGLVQPTRLLLEYLEEKYEEHLYERDEGDKWRNKKFELGLEFPNLPYYI DGDVKLTQSMAIIRYIADKHNMLGGCPKERAEISMLEGAVLDIRYGVSRIAYSKDFETLK VDFLSKLPEMLKMFEDRLCHKTYLNGDHVTHPDFMLYDALDVVLYMDPMCLDAFPKLVCFKKRIEAIPQIDKYLKSSKYIAWPLQGWQATFGGGDHPPKSDLEVLFQGPLGSHLNVLAKALYDNVAESPDELSFRKGDIMTVLEQDTQGLDGWWLCSLHGRQGIVPGNRLKILVG MYDKKP

>BRAF-RBD 155-227GGSPIVRVFLPNKQRTVVPARCGVTVRDSLKKALMMRGLIPECCAVYRIQDGEKKPIGWDTDISWLTGEELHVEVL

### Protein expression and purification

All human and mouse INMT proteins were overexpressed in E. coli BL21 (DE3) and purified using affinity chromatography and size-exclusion chromatography. Briefly, cells were grown at 37°C in TB medium in the presence of 50 μg/mL of kanamycin to an OD of 0.8, cooled to 17°C, induced with 400 μM isopropyl-1-thio-D-galactopyranoside (IPTG), incubated overnight for 20 hr at 17°C, collected by centrifugation, and stored at −80°C. Cell pellets were lysed in buffer A (50 mM HEPES, pH 7.5, 500 mM NaCl, 1 mM Tris(2-carboxyethyl)phosphine (TCEP), 10% glycerol, and 20 mM imidazole) using Microfluidizer (Microfluidics), and the resulting lysate was centrifuged at 30,000g for 40 min. Ni-NTA beads (Qiagen) were mixed with cleared lysate for 45 min and washed with buffer A. Beads were transferred to an FPLC-compatible column, and the bound protein was washed further with buffer A supplemented with 1.5 M NaCl for 20 column volumes and eluted with buffer B (25 mM HEPES, pH 7.5, 500 mM NaCl, 1 mM TCEP, and 400 mM Imidazole). The eluted sample was concentrated and purified further using a Superdex 75 16/600 column (Cytiva) in buffer C (20 mM HEPES, pH 7.5, 200 mM NaCl, and 1 mM TCEP). The fractions containing INMT protein was concentrated to ∼20-40 mg/mL and stored in −80°C.

### Expression and purification of NEDD9 SH3 domain and BRAF RBD

His(6)-SUMO-NEDD9 (1-64) was cloned into a pET21b vector. The construct was transformed into BL21 (DE3) competent E.coli cells (Invitrogen) which were then cultured in TB medium supplemented with 100 μg/mL ampicillin. Protein expression was induced by 0.5 mM isopropyl b-D-1-thiogalactopyranoside (IPTG) when the optical density of cells at 600 nm (OD_600 nm_) reached 0.8. After that the cells were grown at 18°C for 16-20 hours with 180 rpm shaking and collected by ultracentrifugation. The cell pellet was resuspended for lysis in 20 mM HEPES pH 7.4, 150 mM NaCl, 20 mM imidazole pH 7.4 and 0.5 mM TCEP, supplemented with protease inhibitors and lysozyme. The cells were lysed by microfluidizer, and the lysate was clarified by ultracentrifugation followed by filtration. The clarified lysate was passed over a HisTrap column (Cytiva), washed with 20 mM HEPEs pH 7.4, 150 mM NaCl and 0.5 mM TCEP, 10% glycerol and then eluted over a 20-500 mM imidazole gradient. The appropriate fractions were pooled and then His(6)-SUMO tag was cleaved by ULP1 protease. The protein was passed over the HisTrap column to separate the tag and collected from the fractionated flow-through. For the final size-exclusion chromatography step, the protein was passed over a Superdex 26/600 75 pg column (Cytiva) that was equilibrated with the final buffer containing 20 mM HEPES pH 7.4, 150 mM NaCl, 0.5 mM TCEP (for biochemical assays) or 50 mM HEPES pH 8.5, 150 mM NaCl, 0.5 mM TCEP and 10% glycerol (for crystallography). Purified protein was stored at −80°C.

### Production of ^15^N-NEDD9-SH3

For production of isotope-labelled NEDD9 (1-64) protein for NMR studies, the expression proceeded with the above expression vector, starting with an overnight inoculation of transformed E.coli cells into Luria Broth (LB) medium supplemented with 50 μg/mL carbenicillin. Starter cultures were then used to inoculate 1L of M9 minimal medium, prepared with ^15^N-ammonium chloride (Cambridge Isotopes CIL NLM-467). The culture was grown at 37°C until an optical density (OD_600 nm_) of 1 was reached. 1mM isopropyl b-D-1-thiogalactopyranoside (IPTG) was added for induction of protein expression and the culture was grown at 18°C for 18 hours and then cells were harvested. Purification proceeded as described above for unlabeled protein.

### Crystallization

Proteins were mixed with precipitation solutions (200nL+200nL) and crystallized by hanging-drop vapor diffusion at 20°C. Crystals were transferred briefly into crystallization buffer containing 25% glycerol prior to flash-freezing in liquid nitrogen. For Fragment **2**, all the crystallization trials were performed by hanging drop vapor diffusion method. Pure NEDD9-SH3 (1-64) and compound were mixed at 1:4 ratio (500 μM protein: 2 mM compound) and were incubated overnight at 4°C. The mixture was spun down prior to setting up trays to remove any precipitation. Drops were set up by mixing the protein solution with crystallization solution at 2 μL: 2 μL and 2 μL: 3 μL drop ratios and let to equilibrate against 500 μL of well solution at 8°C. Small rod-shaped crystals grew within a week from solutions comprising 600-800 mM (NH_4_)_2_SO_4_ and 100 mM sodium citrate pH 4-4.6. Crystals were flash frozen in liquid nitrogen using 20% glycerol as cryoprotectant. X-ray data were collected at Brookhaven National Lab (NSLSII AMX). Data were indexed and scaled using XDS.^3^

### Determination

Diffraction data were collected at beamline NE-CAT-24ID-E at the Advanced Photon Source (Argonne National Laboratory). Datasets were integrated and scaled using xds or dials package (v.20200417).^4^ Structures were solved by molecular replacement using the program Phaser (v.2.8.3)^5^ using various search models. Iterative manual model building and refinement using Phenix (v.1.20.1_4487)^6,7^ and Coot (v.0.9.7)^8^ led to the final model with statistics shown in the associated table. For the Fragment **2** complex, the structure was solved by molecular replacement using Phaser and our apo structure. Data collection and refinement statistics are reported in Table S3.

### Fluorescence polarization

Binding assays were performed in 33ul volume with labeled probe at a concentration of 100 nM, the IRF4 protein at 20uM concentrations, and small molecules or peptides ligands titrating within 100nM and 100uM in buffer containing 20 mM Hepes pH 7.6, 20 mM NaCl, 100 uM TCEP, 0.01% Pluronic acid. FP assays were performed in 384 well plates and read using a PherastarFS plate reader.

### Isothermal titration calorimetry

All ITC measurements were carried out at 25°C using an Affinity ITC (TA Instruments, Waters) equipped with auto sampler. 20uM NEDD9 solution (50mM Hepes-7.5, 150mM NaCl, 2% DMSO) in the calorimetric cell was titrated with 2.5uL injections of 200uM ligand solutions (50mM Hepes-7.5, 150mM NaCl, 2% DMSO) using 200 sec intervals and stirring speed at 125rpm. Resulting isotherm was fitted with a single site model to yield thermodynamic parameters of H, S, K_D_, and stoichiometry using NanoAnalyze software (TA instruments). Calorimetric data was collected using an AffinityITC Calorimeter provided by TA-Instruments.

### Flow-induced dispersion analysis

FIDA was carried out in the FIDA1 instrument (FIDA Biosystems ApS, Denmark) using the standard protocol recommended by the manufacturer.^9^ In short, NEDD9 SH3 (55 uM) was titrated with compound solution at various concentrations (0 - 300 uM) by injecting protein solution into a stream of compound solution and allowing binding to take place during the run. The complex formation could then be observed using intrinsic fluorescence of NEDD9. The resulting hydrodynamic radius was calculated for each concentration of the compound, and K_D_ was calculated by fitting the binding curve to a 1:1 model using the FIDA software 2.3 (FIDA Biosystems ApS, Denmark). In triplicate, the analyses were performed in a PBS buffer, pH 7.4, containing 5% DMSO.

### Intact mass analysis by LC/MS

Recombinant NEDD9-SH3 domain was analyzed at 1 uM, with compounds and alkylating reagents at 100 μM unless otherwise indicated within figure legends. Incubations were carried out at 21°C for one hour unless otherwise indicated in reaction buffer containing 50 mM HEPES, pH 7.4 and 100 mM NaCl. Reactions were quenched with formic acid to a final concentration of 0.2% and analyzed using the Waters Bioaccord LC-TOF. Mobile phase A consisted of 0.1% formic acid in LC-MS grade water and mobile phase B consisted of 0.1% formic acid in LC-MS grade acetonitrile. Protein was injected onto a reverse-phase C4 column (ACQUITY UPLC Protein BEH, 300Å, 1.7 µm, 2.1 × 50 mm) held at 80 °C and desalted for one minute before elution with a gradient of 5% to 85% acetonitrile in 2.5 min. Electrospray ionization (ESI) was used in positive mode with the cone voltage set to 55 V and 550 °C desolvation temperature. Protein mass was deconvoluted using MaxEnt1 in Waters UNIFI software.

### 2D NMR experiments with 15N-NEDD9

2D HSQC experiments with ^15^N-NEDD9 were performed on a Bruker Avance 600 MHz NMR spectrometer equipped with a cryoprobe. Samples contained 75uM 15N-NEDD9 with or without compound in buffer containing 20mM Hepes-d_16_, pH 7.4, 150mM NaCl, prepared with 10% D_2_O. Compounds were added at a final concentration of 200 μM (Fragment **1**) or 150 μM (Fragment **2**) with 2% DMSO-d_6_. For Fragment **2**, 0.5 mM TCEP-d_11_ was included in the buffer. Deuterated reagents were obtained from Cambridge Isotope Laboratories, Inc. HSQC experiments were acquired with 64 scans at a temperature of 298K using a Fast-HSQC pulse program for improved sensitivity.^10,11^

### Covalent library screening by RapidFire-MS and kinetic analysis

We screened N=6,120 covalent electrophiles (Enamine®) against the NEDD9-SH3 domain by RapidFire-MS (Momentum Biotechnologies). Equimolar amounts of NEDD9-SH3 and the BRAF-Ras binding domain (RBD) in the same reaction were incubated with the fragment library as a means of performing the screen and counter screen simultaneously. 400 nL of 10 mM stock of the covalent fragment library was dispensed using the Echo Liquid Handler (Beckman Coulter). In a final volume of 40 μL, 1 μM NEDD9-SH3 and 1 μM BRAF(RBD) were incubated in the presence of 100 μM compound, 100 μM iodoacetamide or 1% DMSO in 50 mM HEPES, pH 7.4, 100 mM NaCl for 4h at 21°C. Reactions were quenched with formic acid for a final concentration of 0.2%. Plates were sealed, frozen at −80°C and shipped on dry ice to Momentum Biotechnologies (Billerica, MA) for RapidFire-MS analysis on the Agilent QTOF system. Compounds that formed a single adduct to NEDD9-SH3 at least two-fold greater than to BRAF-RBD were considered hits and moved forward in the screening funnel. Further kinetic analysis was performed by varying the compound concentration and incubation time as shown in the figures. Analysis of these reactions was performed as described above in “*Intact Mass analysis by LC/MS*”.

### Kinetic evaluation of compound occupancy by mass spectrometry

The covalent inhibition kinetics of BRD-K00098212-001-02-9 against the SH3 domain of NEDD9 were assessed by time-dependent occupancy measurements using intact protein mass spectrometry. The SH3 domain (1 μM final concentration) was incubated with six concentrations of compound (4.7, 9.4, 18.7, 37.5, 75, and 150 μM) in 2% DMSO at 37 °C. Reactions were quenched at 0.25, 0.5, 1, 2, 4, and 6 hours by addition of 2% formic acid. Ubiquitin (100 nM) was included as an internal standard. A 384-well plate format was used with a reaction volume of 25 µL transferred into quench plates for each timepoint. Samples were analyzed on a Thermo Scientific Q Exactive HF (QE-HF) Orbitrap mass spectrometer to detect covalently modified NEDD9. Peak deconvolution was performed using Biopharma Finder, and occupancy data were fitted to determine kinetic parameters using a custom Python script.

### Peptide mapping

To identify cysteine residues on NEDD9 modified by Fragment 2, full-length NEDD9 or its SH3 domain (2 μM) was incubated with compound (typically 100 μM final) for 30 minutes at room temperature. Following the reaction, protein was denatured using 2× lysis buffer (10% SDS, 100 mM TEAB, pH 8.5), reduced with 5 mM TCEP, and alkylated with 20 mM iodoacetamide (IAA) in the dark for 30 minutes. Samples were acidified with 2.5 μL of 27.5% phosphoric acid to achieve a final pH ≤ 1 and loaded onto S-Trap™ micro spin columns (ProtiFi, LLC). Protein digestion was carried out on-column with trypsin (1:10 enzyme-to-substrate ratio) in 50 mM TEAB buffer at 47 °C for 1.5 hours.

Peptides were sequentially eluted in three fractions using (i) 50 mM TEAB, (ii) 0.2% formic acid in water, and (iii) 50% acetonitrile in water, each at 40 μL, and then dried by vacuum centrifugation. Reconstituted peptides were analyzed by LC-MS/MS using a Q-Exactive HF Orbitrap mass spectrometer connected to a Vanquish Horizon UHPLC system (Thermo Fisher Scientific, Bremen, Germany). All experiments were conducted in positive electrospray ionization (ESI) mode. Peptide mapping data were processed with Proteome Discoverer (Thermo Fisher Scientific) and manual MS/MS inspection to confirm covalent labeling. The resulting spectra allowed identification of Cys18 as the primary labeled residue.

### Computational folding

Structure prediction by Alphafold.^12^

### Peptide synthesis

Enter motif (PxKPxR query pattern) in https://www.genome.jp/tools/motif/MOTIF2.html searching the Swiss-Prot database to produce a dataset. Peptides corresponding to amino acids 678-686 of human FAK1 containing bromo acetamide side chain variants were generated using solid-phase Fmoc chemistry on a Symphony X peptide synthesizer (Gyros Protein Technologies), with amino acids sequentially added to rink amide AM resin. Following N-terminal acetylation, deprotection, and cleavage from the resin, peptides were purified by reverse-phase high performance liquid chromatography and mass spectrometry (LC-MS) and quantified by amino acid analysis.

### Angiogenesis assay

Endothelial tube formation was analyzed on glass chamber slides (Lab-Tek II) covered with extracellular matrix gel (Abcam, catalog #ab204726) according to manufacturer recommendations. HPAECs or PAH-HPAECs were incubated on extracellular matrix-coated slides for 6 hours at 37°C, 5.0% CO_2_, fixed with 4% phosphate-buffered formalin and images were taken using Leica DMi8 inverted microscope. Branch points were counted in 3-4 random 10x magnification view-fields per well. To analyze the effect of Sulf1 overexpression or inhibition on tube formation, HPAECs were incubated for 24 hours with adenovirus expressing GFP (Ad-GFP), GFP plus human Sulfatase-1 (Ad-Sulf1) or GFP plus human Sulf1-siRNA (Ad-siSulf1), 1 PFU/ml x10^7^ (Vector Biolabs) before plating on extracellular matrix-coated slides.

### Wound healing assay

HPAECs cultured on glass chamber slides (Lab-Tek II) were scratched using 100 μl pipette tip and fixed with 4% formaldehyde 6 hours later. Average distance between migrating cells at three different points was measured on 10x magnification images.

### Sulf1 inhibition in HPAECs

Cells were transfected using Human Artery Endothelial Cell Avalanche Transfection Reagent (EZBiosystem, catalog# EZT-HAOE-1) according to manufacturer recommendations. siRNA from Ambion was used: sense sequence – CGGAUACAGCAGGAACGAAUU, antisense sequence – 5’-P-UUCGUUCCUGCUGUAUCCGUC. Control cells were treated with control siRNA-A (Santa Cruz, catalog# sc-37007). Cells were used for wound healing assay 24 hours after transfection.

### Immunoblotting

Proteins were size-fractionated electrophoretically using sodium dodecyl sulfate polyacrylamide gel (SDS-PAGE) and transferred to polyvinylidene fluoride membranes according to methods reported previously.^2^ The membranes were incubated with anti-NEDD9 (Abcam, catalog# ab18056) [dilution 1:1000], SMAD3 (Abcam, catalog# ab40854) [dilution 1:1000], anti-GBR2 (Invitrogen catalog# PA5-27151) [dilution1:500] overnight at 4°C, incubated with peroxidase-labeled secondary antibody, and visualized using the ECL detection system (Amersham Biosciences).

### Co-immunoprecipitation

Co-immunoprecipitation was performed on HPAECs monolayers as reported previously^2^ using SureBeads Protein G magnetic beads (Biorad) and NEDD9 antibody (Abcam, catalog# 18056) according to manufacturer protocol.

### Actin staining

HPAECs were fixed with 4% formaldehyde and stained using actin cytoskeleton and focal adhesion staining kit (Millipore, catalog# FAK100) according to manufacturer protocol.

### NEDDtide transfection

Wound healing assay. HPAECs were transfected with the relevant bromoacetylated peptides using Chariot^TM^ Protein Delivery Reagent (Active Motif, catalog# 30025) according to manufacturer protocol. After scratching, HPAECs were incubated in serum-free medium containing Chariot-peptide complexes (5 μM) for 3 hr, and then equal amount of serum-containing medium was added for another 3 hours until fixation and analysis.

### Immunoblot

HPAECs were grown to 80% confluence using complete endothelial basal medium-2 from Lonza (catalog # cc3156). HPAECs were transfected with the transfection reagent Chariot from Active Motif, with the different peptides, and the positive control β-galactosidase using the manufacturer’s protocol with modifications. Proteins were isolated immediately in the presence of protease inhibitor cocktail (catalog# 539134) after the transfection experiments for immunoblot. Proteins were electrophoresed in a TGX stain-free polyacrylamide gel (catalog # 4568094), using Mini PROTEIN tetra cell from Bio-Rad, and transferred to 0.2 μm PVDF membrane by Trans Blot Turbo transfer system using semi-dry rapid blotting technology. The membranes were then blocked with 5% non-fat milk to avoid non-specific binding and incubated with an anti-NEDD9 antibody (Abcam, catalog # ab18056; [dilution 1:1000]), gel loading control anti-beta-actin (Cell Signaling, catalog # 4967: [1:1000]) or GAPDH (Millipore, catalog # G8795), and another antibody for non-canonical protein of the human SH3 domain family, anti-GBR2 (Invitrogen PA5-27151: [1:500]) at 4C for overnight. Proteins were visualized by enhanced chemiluminescence substrate (Western Bright^TM^ ECL from Advansta; catalog # K-12045-D50) marked by the respective secondary antibodies’ horseradish peroxidase (HRP).

### Immunofluorescence in vitro

HPAECs were grown on glass chamber slides (Lab-Tek II), fixed with 4% formalin in PBS, blocked with 10% goat albumin (Life Technologies) in phosphate-buffered saline (PBS) for 1 hr at room temperature and incubated with primary antibody (dilution 1:100) overnight at 4°C. After washing cells were incubated for 1 hour with secondary antibodies: goat anti-rabbit conjugated with Alexa Fluor 568 (Abcam catalog# 175471) and/or goat anti-mouse conjugated with Alexa Fluor 488 (Abcam catalog# 150113) and mounted on glass slides with anti-fade mounting medium with DAPI (Vector Laboratories).

### Animal experiments; general methodology

Animals were selected randomly for experimentation, and data analysis was performed using a blinded methodology. All animals were included in experiments, and only animals that endured substantial bleeding, respiratory failure during anesthesia, or died from a complication of the catheterization procedure (described below) were excluded from analysis.

### Histology and immunohistochemistry

Animal lung samples were perfused with 10% phosphate-buffered formalin at a pressure of 20 cm H_2_O prior to harvesting and then fixed with formalin for at least 24 hours at room temperature.^2^ Samples were processed/embedded in paraffin using a Hypercenter XP System and Embedding Center (Shandon). The paraffin-embedded lung tissue was cut into 5-µm sections and used for histological and immunohistochemical analysis. To assess overall collagen deposition and fibrillar collagen in specific, sections were stained with a Gomori’s Trichrome Staining kit (Fisher Scientific) and Picrosirius Red Stain Kit, respectively, according to manufacturer’s instructions. Vascular fibrosis was analyzed on vessels with an approximate diameter of 20–50 µm, located distal to terminal bronchioles The degree of vascular remodeling was calculated by subtracting the area of the lesser curvature from the area of the greater curvature and dividing by the greater curvature x 100, as reported previously.^13^ For immunohistochemical analysis lung sections were blocked with 10% goat albumin (Life Technologies) in phosphate-buffered saline (PBS) for 1 hour at room temperature and incubated with primary antibody (dilution 1:100) overnight at 4°C. After washing cells were incubated for 1 hour with secondary antibodies: Alexa Fluor 568 (Abcam catalog# 175471) and/or goat anti-mouse conjugated with Alexa Fluor 488 (Abcam catalog# 150113) and mounted on glass slides with anti-fade mounting medium with DAPI (Vector Laboratories). Images were acquired using an LSM 700 Flexible Confocal Microscope (Carl Zeiss) and Leica DMi8 inverted microscope and analyzed using Zen and Leica LAS X software correspondingly.

### Endothelial collagen synthesis in vitro

HPAECs were grown to 50% confluence in chamber slides and transfected with Sulf1 adenovirus expressing GFP plus human Sulfatase-1 (Ad-Sulf1) or GFP (Ad-GFP) as a control, 1 PFU/ml x10^7^ (Vector Biolabs). Three days after transfection cells were fixed and stained with Collagen III antibody (Abcam, catalog# 7778).

### Podosome detection

HPAECs were grown on glass chamber slides (Lab-Tek II) and transfected for 24 hours with adenovirus expressing GFP (Ad-GFP), GFP plus human Sulfatase-1 (Ad-Sulf1) or GFP plus human Sulf1 siRNA (Ad-siSulf1), 1 PFU/ml x10^7^ (Vector Biolabs) and stained with TRITC-conjugated phalloidin for actin and following antibodies: phospho-FAK (Tyr861), phospho-FAK (Tyr576), vinculin, cortactin, sulfatase-1 and neuropilin-1. The rate of podosome formation was measured as a percentage of cells expressing phospho-FAK (Tyr861)-positive circles and crescents counted per view field on 20x magnification images.

### Immunogold Electron Microscopy

HPAECs and HPASMCs were co-cultured on glass chamber slides (Lab-Tek II) and fixed with 2.5% Glutaraldehyde 2.5% Paraformaldehyde in 0.1 M sodium cacodylate buffer (pH 7.4) for 1 hour at RT. After blocking with 10%, 1% BSA in PBS for 1 hour at RT cells were incubated with Neuropilin-1 (1:30 dilution) in 1% BSA in phosphate-buffered saline overnight at 4°C. After washing in PBS, cells were incubated with Protein A gold-15 nm (University Medical Center Utrecht, The Netherland) in 1% BSA in PBS for 30 min at RT. After washing in PBS cells were fixed for 1 minute in 0.5% glutaraldehyde, washed 4 x in 20mM Glycine/PBS to quench free aldehyde groups and incubated with Heparan Sulfate antibody (1:20 dilution) in 1% BSA in PBS overnight at 4°C. After washing in PBS, cells were incubated with Protein A gold-10 nm (University Medical Center Utrecht, The Netherland) in 1% BSA in PBS for 30 min at RT. After washing in PBS cells were fixed for 1 minute in 0.5% glutaraldehyde and then washed 2x in 0.1M Sodium Cacodylate buffer pH 7.4, postfixed in 1% Osmium tetroxide (OsO4)/1.5% Potassium ferrocyanide (KFeCN6) for 30 min, washed 2x in water, 1x Maleate buffer (MB) and incubated in 1% uranyl acetate in MB for 30 min followed by 1 wash in MB and 2 washes in water and subsequent dehydration in grades of alcohol (5min each; 50%, 70%, 95%, 2x 100%). Cells were embedded in Epon-Araldite resin, trimmed to 0.5×1 mm rectangle and cut into ultrathin sections (about 60nm) on a Reichert Ultracut-S microtome and examine in the JEOL 1200EX Transmission electron microscope with an AMT 2k CCD camera.

### Animal models of PAH

#### Monocrotaline rats

As previously described,^2^ male Sprague-Dawley rats (12-14 weeks of age; Charles River Laboratory) were administered a single intraperitoneal injection of monocrotaline (60 mg/kg) (Sigma) and experiments were performed 23 days later.

#### Sugen-5416-Hypoxia-Normoxia Rats

Male Sprague-Dawley rats (12-14 weeks of age; Charles River Laboratory) were administered a single subcutaneous injection of the VEGFR-2 kinase inhibitor Sugen-5416 (20 mg/kg; Sigma), exposed immediately to chronic hypoxia (10% FiO_2_) for 21 d, and then returned to normoxia until completion of the protocol 32 days later, as reported previously.^2^

#### Sugen-5416-Hypoxia-Normoxia PAH reversal treatment protocol with NEDD9 siRNA administration

Intratracheal injection of NEDD9 or control non-targeting si-RNA (1mg/kg of body weight) began on day 14 following initiation of SU-5416-Hypoxia treatment and was repeated on day 19 and 25. ON-TARGETplus Rat NEDD9 siRNA from Dharmacon was used; sense sequence – CAAUGAAACAAGACGGAAAUU, antisense sequence – 5’-P-UUUCCGUCUUGUUUCAUUGUU. Control group was treated with ON-TARGETplus Non-targeting siRNA; sense sequence – UGGUUUACAUGUCGACUAAUU, antisense sequence – 5’-P-UUAGUCGACAUGUAAACCAUU. As a transfection agent Avalanche-in vivo Transfection Reagent (EZ Biosystems^TM^ Cat. No. EZT-VIVO-1) was used according to manufacturer recommendations.

#### Monocrotaline and Sugen-5416-Hypoxia-Normoxia PAH prevention treatment protocols with Sulf1 siRNA administration

In monocrotaline model intratracheal injection of Sulf1 si-RNA or control non-targeting si-RNA (1mg/kg of body weight) was performed on days 1, 8 and 15 and in Sugen-5416/Hypoxia-Normoxia model on days 7, 16 and 23. ON-TARGET plus Rat Sulf1 siRNA-SMART pool was used (Dharmacon Cat. No. L-093746-02-0050); target sequence 1: GGUAACAGGUUUCGAACAA, target sequence 2: GGAAGUACGUGCAAUAACCA, target sequence 3: CACAAGGCCUACAUCGAUA, target sequence 4: CCAACGUCCUCCAGCGCAA. Control group was treated with ON-TARGETplus Non-targeting siRNA; sense sequence – UGGUUUACAUGUCGACUAAUU, antisense sequence – 5’-P-UUAGUCGACAUGUAAACCAUU. As a transfection agent Avalanche-in vivo Transfection Reagent (EZ Biosystems^TM^ Cat. No. EZT-VIVO-1) was used according to manufacturer recommendations.

#### Hemodynamic analyses

All experiments were performed with rats sedated using ketamine (50 mg/kg) and xylazine (10 mg/kg) according to methods published by our laboratory previously.^2,14^

#### Rat right heart catheterization and hemodynamics

A 2 F high fidelity Millar conductance catheter (Millar Instruments, Inc.; Model SPR-869) was inserted into the carotid artery; the distal clamp was then released, and the suture was fastened sufficiently to prevent substantial blood loss. The catheter was advanced into the aorta and left ventricle (LV), and aortic mean arterial pressure, cardiac output, and left ventricular end-diastolic pressure was recorded at steady-state (digitally recorded by MPVS-400, Power Lab 8/35, 1 kHz sampling rate; Millar Instruments and ADInstruments, Inc.). Right ventricular systolic and end-diastolic pressures were measured by fluid-filled catheter as previously described.^2^ Data were analyzed in Lab Chart Pro v8.1.5 (ADInstruments, Inc.).

#### Right ventricular weight

The RV and the LV+septum weight (Fulton index) were recorded as reported previously.^2^ Data were expressed as the ratio of RV weight (g)/LV + septum weight (g).

#### Statistical methods

All analyses were performed using Origin 2025^TM^. Data are expressed as mean ± S.E.M. Comparison between two groups was performed by the Student’s unpaired two-tailed t-test. One-way analysis of variance (ANOVA) was used to examine differences in response to treatments between >2 groups. Post-hoc analysis was performed by the method of Tukey. P<0.05 was considered significant.

## Acknowledgements

The authors wish to acknowledge the contributions of Puspalata Bashyal, M.Sc., Alex Burgin, Ph.D., Maxime Descourt, M.S., Israa El Saudi, M.Sc., Maria Ericsson, B.Sc., Carla M. Gauss, Ph.D., Ezekiel Geffken, M.Sc., Alexandra Gould, Ph.D., Hsueh-Cheng Huang, Ph.D., William Kubasek, Ph.D., Enrico Margiotta, Ph.D., Aroonroj Mekareeya, Ph.D., Coby O’young, B.S., Valeria Padovano, Ph.D., Luke Sebastian, B.S., Mrinal Shekhar, Ph.D., Andrew W. Stamford, Ph.D., Nick Vangos, B.S., Zoe Yeoh, B.S. This work was based upon research conducted at the Northeastern Collaborative Access Team beamlines, which were funded by the National Institute of General Medical Sciences from the National Institutes of Health (P30 GM124165). The Eiger 16M detector on 24-ID-E was funded by a NIH-ORIP HEI grant (S10OD021527). This research used resources of the Advanced Photon Source, a U.S. Department of Energy (DOE) Office of Science User Facility operated for the DOE Office of Science by Argonne National Laboratory under Contract No. DE-AC02-06CH11357. BNL-17ID-1/2: The Center for Bio-Molecular Structure (CBMS) is primarily supported by the NIH-NIGMS through a Center Core P30 Grant (P30GM133893), and by the DOE Office of Biological and Environmental Research (KP1607011). NSLS2 is a U.S.DOE Office of Science User Facility operated under Contract No. DE-SC0012704. BNL-17ID-1/2 via APS (BAG-311950).

## Author Contributions

A.O.S., conceptual design, *in vitro*, *in cellulo*, *in vivo* experimental planning and completion, manuscript drafting, manuscript editing; H-S. S., crystallography, *in vitro* chemical assays; A.L., 2-D NMR, project leadership, manuscript editing; B.H., x-ray crystallography, manuscript editing; G.H.B., peptide synthesis and conceptual contributions; D.M., chemistry experiment planning and completion, manuscript edting; P.S., *in cellulo* experimental planning and completion; J.D.; *in vivo* experimental planning and completion; J.M.; mass spectrometry assay development, project leadership; J.Y., 2-D NMR; D.K., chemistry experiment planning and completion and project leadership; W.K., conceptual design and team leadership; J.K., chemistry assay design and synthesis; N.L., mass spectrometry data analysis and assay development; A.M., screening cascade and chemistry assay development; A.B., conceptual design; A.G.; conceptual design; V.K., conceptual design and team leadership; L.D.W., conceptual contributions, manuscript review; P.C., chemistry assay design and synthesis; W.M.O., conceptual design, manuscript editing; M.L.S., conceptual design and manuscript editing; S. D-P., conceptual design, protein synthesis, x-ray crystallography, chemical assay design and completion; B.A.M., conceptual design, overall experimental planning, manuscript drafting, manuscript editing, funding.

## Supplemental Figure Legends

**Figure S1.**
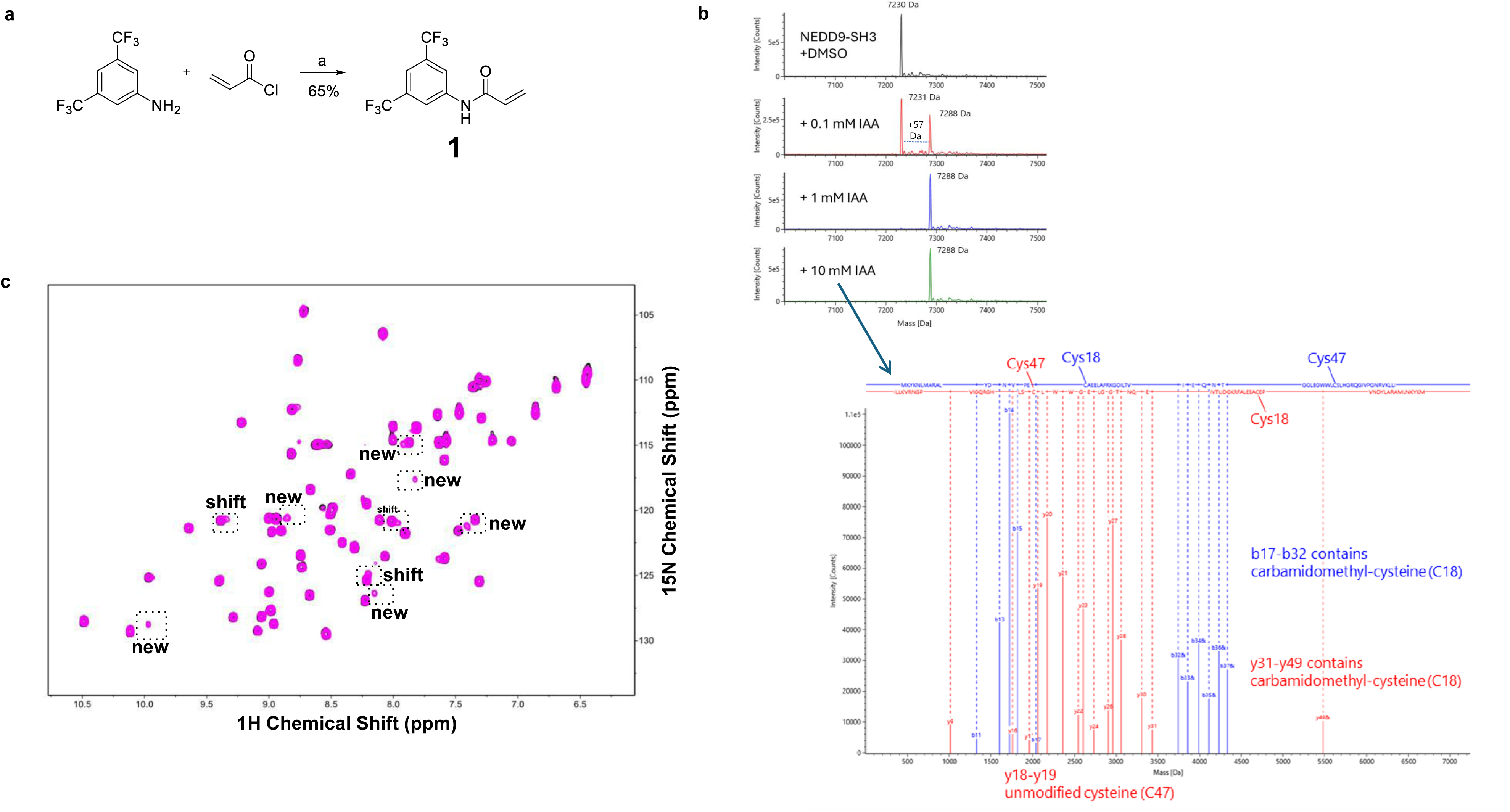
Biochemical profile of Fragment 1. **a.** Synthesis of *N*-(3,5-Bis[trifluoromethyl]phenyl) acrylamide **1**. ^a^Reagents and conditions: (a) acryloyl chloride, DIPEA, CH_2_Cl_2_, 0 °C, 2 h. **b.** The deconvoluted, intact average mass spectra of NEDD9-SH3 protein modified by increasing concentrations of iodoacetamide, resulting in a mass shift of 57 Da (top). MS2 fragmentation showing the protein is only modified on Cys18 and not Cys47 with b-ions in blue and y-ions in red (bottom). **c.** 2D 15N/1H TROSY-HSQC spectra of 75 µM 15N-NEDD9 (1-64) alone in black and in the presence of 200µM **1** in magenta. New amide peaks and shifted peaks in the presence of **1** are indicated. ppm, parts per million.

**Fig. S2.**
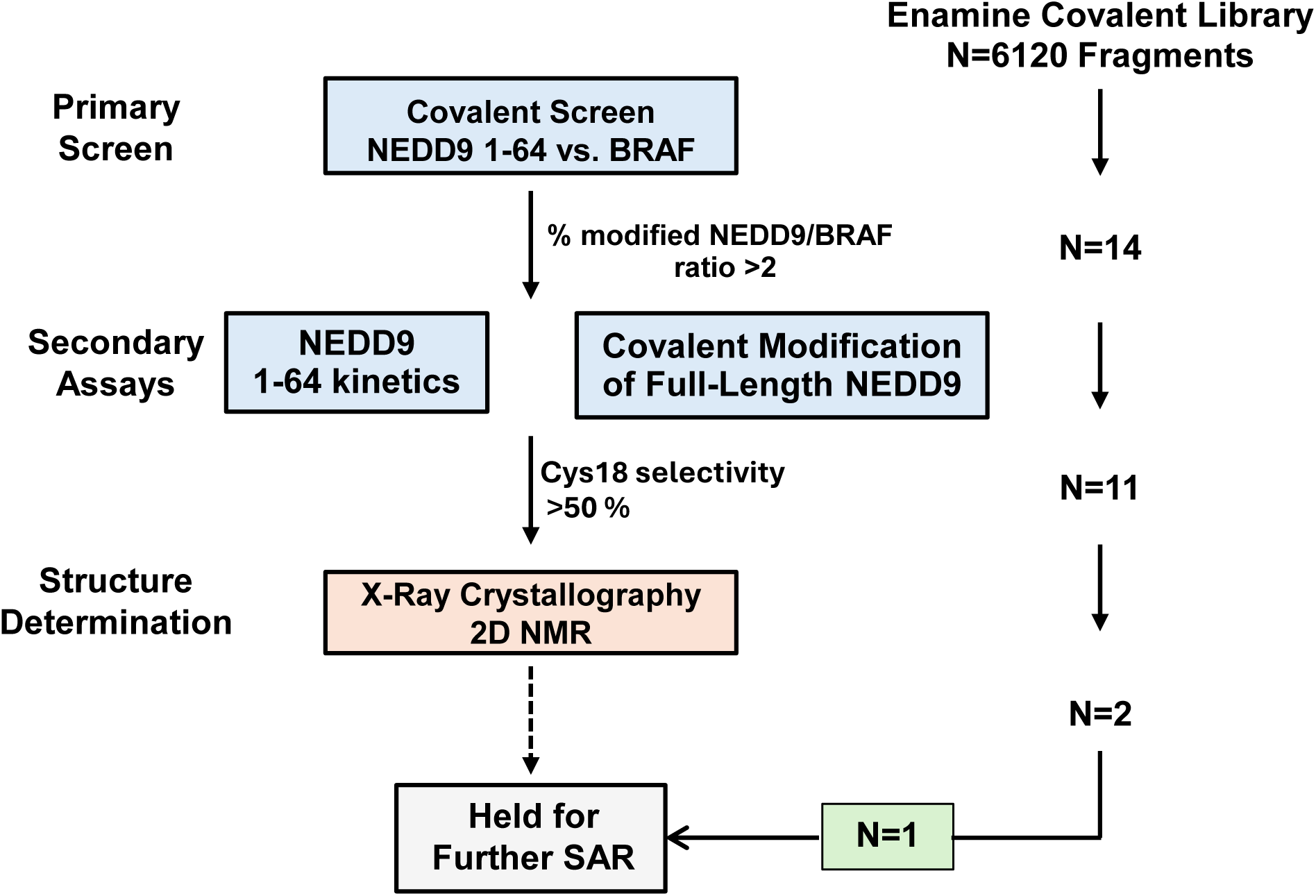
Throughput and results of a small molecule screening funnel for NEDD9-Cys18 interactors. Overall approach to the screening funnel. To remove non-selective alkylating agents early in the screening, BRAF-RBD was included as a counter screen. This was chosen due to a similarly solvent-exposed cysteine compared to NEDD9-SH3 and similar but non-overlapping mass-to-charge peaks in the TOF-MS analysis, allowing both proteins and their adducts to be assayed simultaneously. Light blue, mass spectrometry; NMR, nuclear magnetic resonance; SAR, structure activity relationship.

**Fig. S3.**
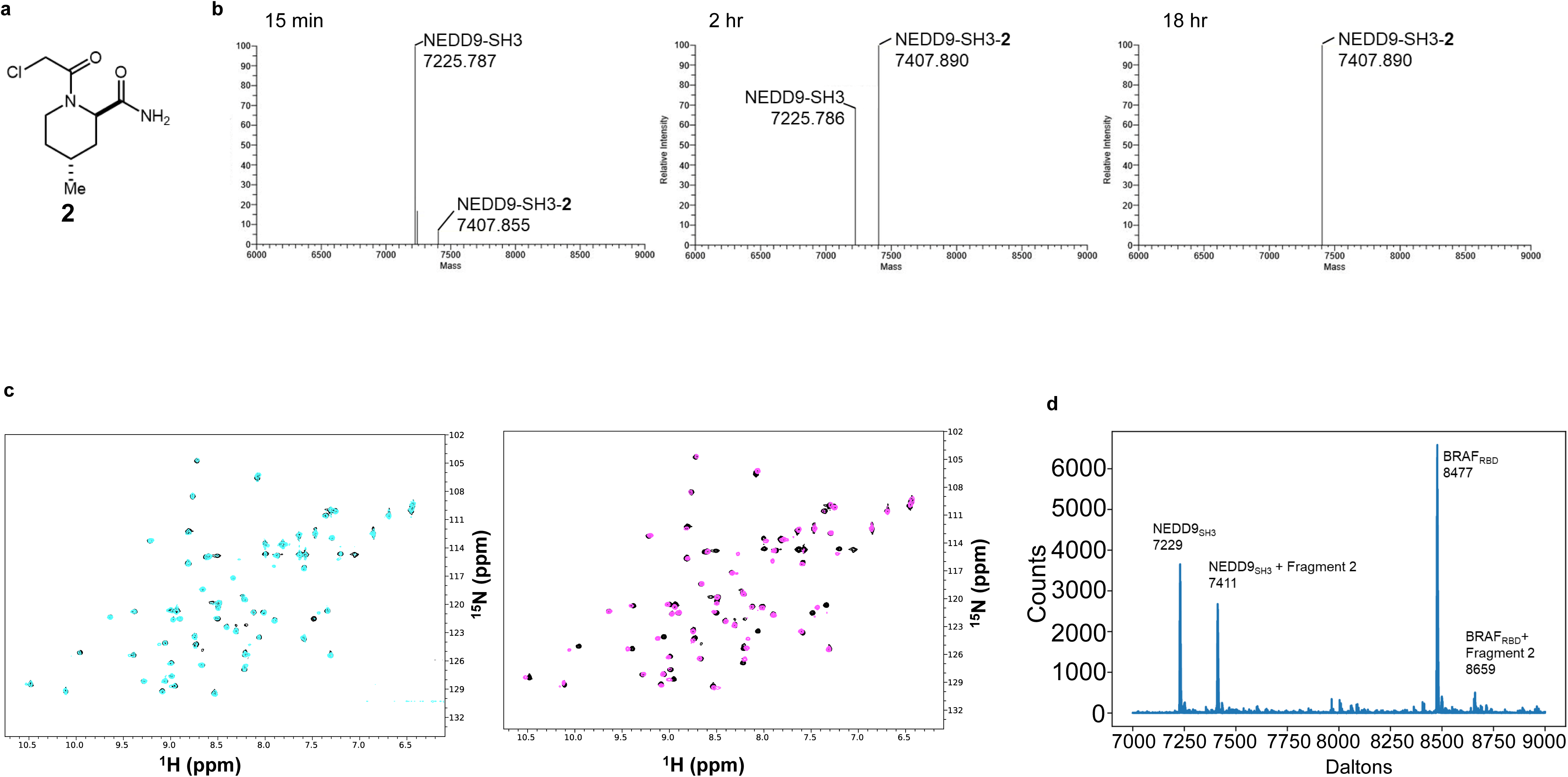
Structure and NEDD9-Cys18 interacting profile of Fragment 2. **a.** Chemical structure of **2. b.** The NEDD9-SH3 construct was incubated with **2** for 15 min, 2 hr, and 18 hr and analyzed by intact MS, which confirmed a time-dependent increase in Cys18 modification by **2** (*n*≥3). The same treatment results in no modification of BRAF-RBD, used as a counterscreen. The deconvoluted monoisotopic masses are shown. **c.** 2-D 15N/1H TROSY-HSQC spectra of 75 µM 15N-NEDD9 (1-64) alone in black and in the presence of 150µM **2** at timepoint 0 hr at room temperature in cyan and after 24 hr in magenta. **d.** Deconvoluted MS spectrum showing NEDD9, NEDD9 modified with fragment 2 (Ratio 56:42), BRAF (calc 8477), and BRAF modified with Fragment 2 (Ratio 94:6) after 4 hr incubation.

**Fig. S4.**
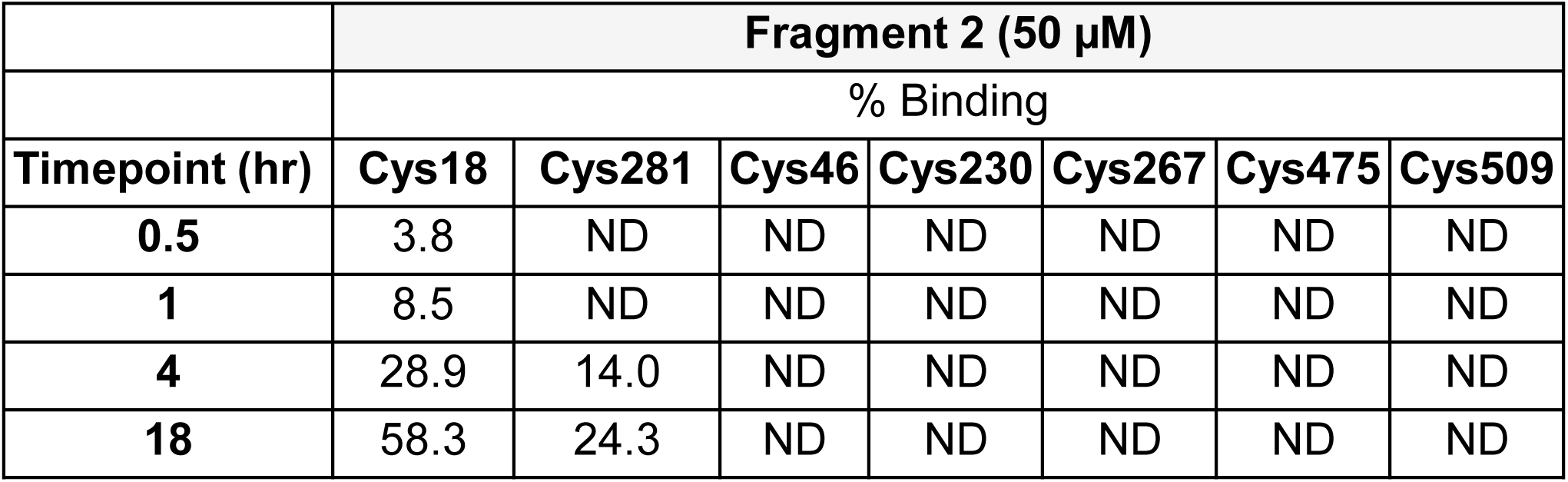
Interaction between Fragment 2 and selected NEDD9 cysteines. The full length NEDD9 construct was incubated with 2 for various time points and the cysteine occupancy profile was reported by liquid chromatography-mass spectrometry on a Q-Exactive HF mass spectrometer (*n*≥3 per cysteine analysis). ND, not detected.

**Fig. S5.**
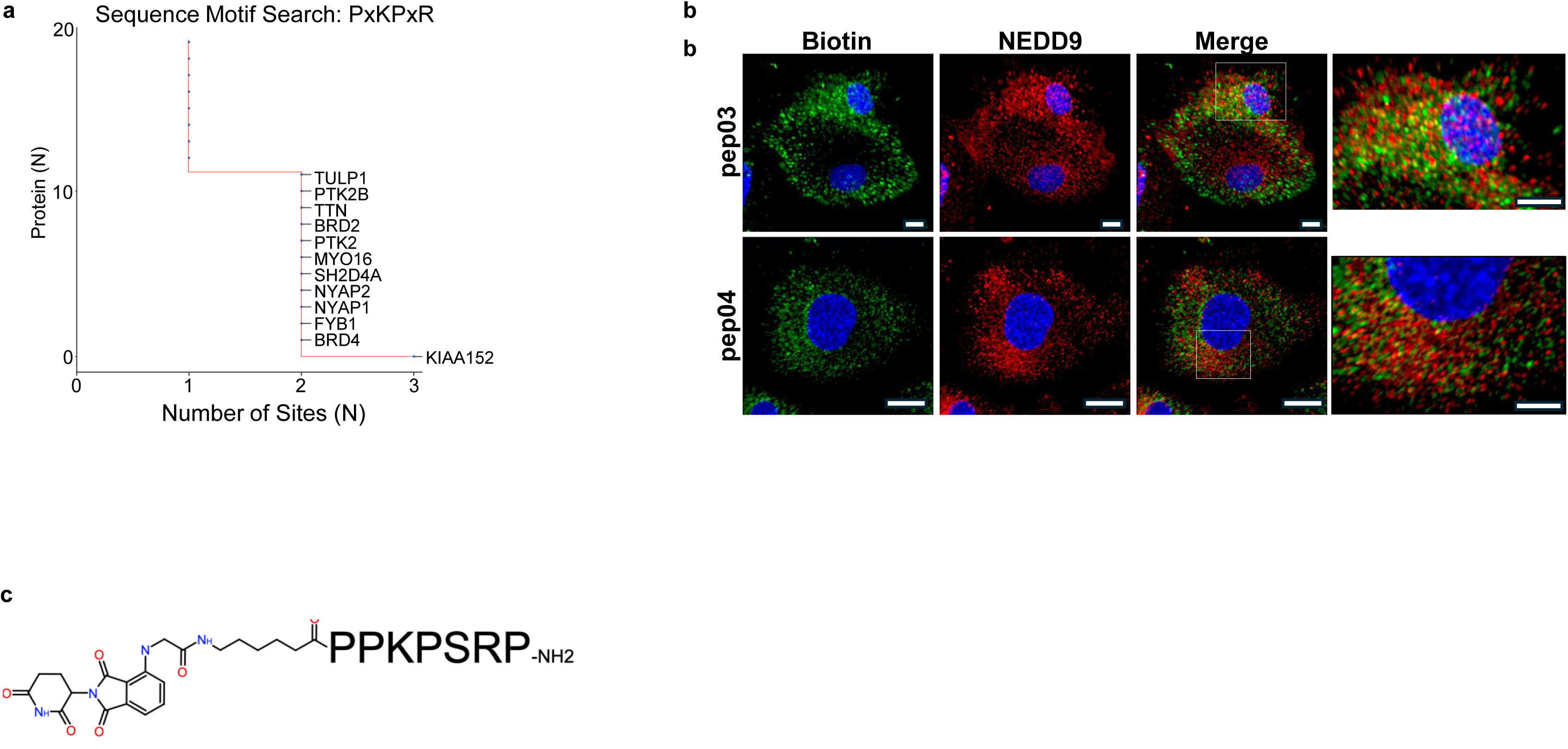
NEDDtide development and validation. **a.** Results of a bioinformatics analysis profiling proteins in the proteome that express the PxKPxP consensus motif. Proteome sequence analysis using Motif2^26^ revealed human genes with three, two and one occurrences of the motif, including FAK1 and FAK2. **b.** Human pulmonary artery endothelial cells were incubated with NEDDtide pep01 modified with a biotin tag (which is pep03) or a scrambled control (Scr) peptide for 2 hr. Anti-biotin (red) and anti-NEDD9 (green) immunofluorescence was performed. Blue, Dapi. The results of this analysis demonstrated biotin-NEDD9 colocalization (yellow) at subcellular structures resembling focal adhesions in pep03-transfected but not Scr-transfected cells (*n*=3). Scale bar, 20µm. **c.** Structure of NEDDtide pep05 including thalidomide moiety.

**Fig. S6.**
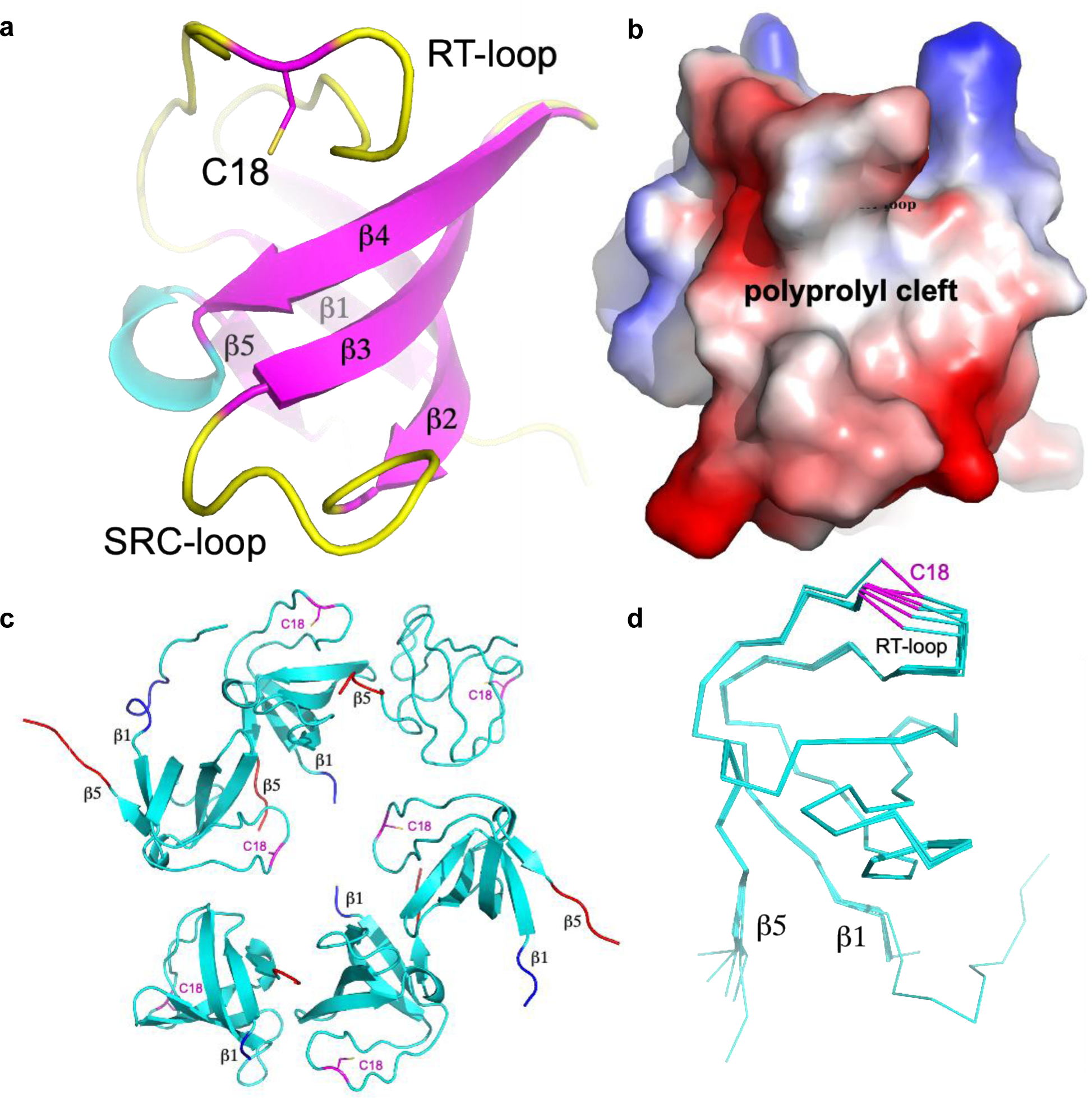
Structural Features of the Apo NEDD9 SH3 domain highlighting the RT-loop and C18 residue. **a.** Cartoon representation of the NEDD9 domain showing the secondary structure elements including five beta-strands (beta1-beta5, magenta) forming the core beta-sheet. The RT-loop, a flexible region critical for substrate binding, is highlighted in yellow. The side chain of residue C18 is shown in stick representation. An inset displays the local conformation around C18. **b.** Electrostatic surface representation of the NEDD9 domain illustrating the proline-rich cleft (labeled) where substrate binding typically occurs. The RT-loop and C18 are positioned adjacent to this cleft, suggesting a role in modulating access or substrate orientation. **c.** Structural alignment of multiple NEDD9 homologs (cyan) to illustrate conservation of global fold. The RT-loops (pink), beta1 (purple), and beta5 (red) strands are annotated, and the positions of C18 in each structure are marked. **d.** Superposition of NEDD9 domain backbone traces highlighting structural variability in the RT-loop region. The beta1 and beta5 strands are labeled for orientation. The RT-loop and C18 side chain (magenta) show variable positioning among structures, emphasizing potential functional flexibility.

**Fig. S7.**
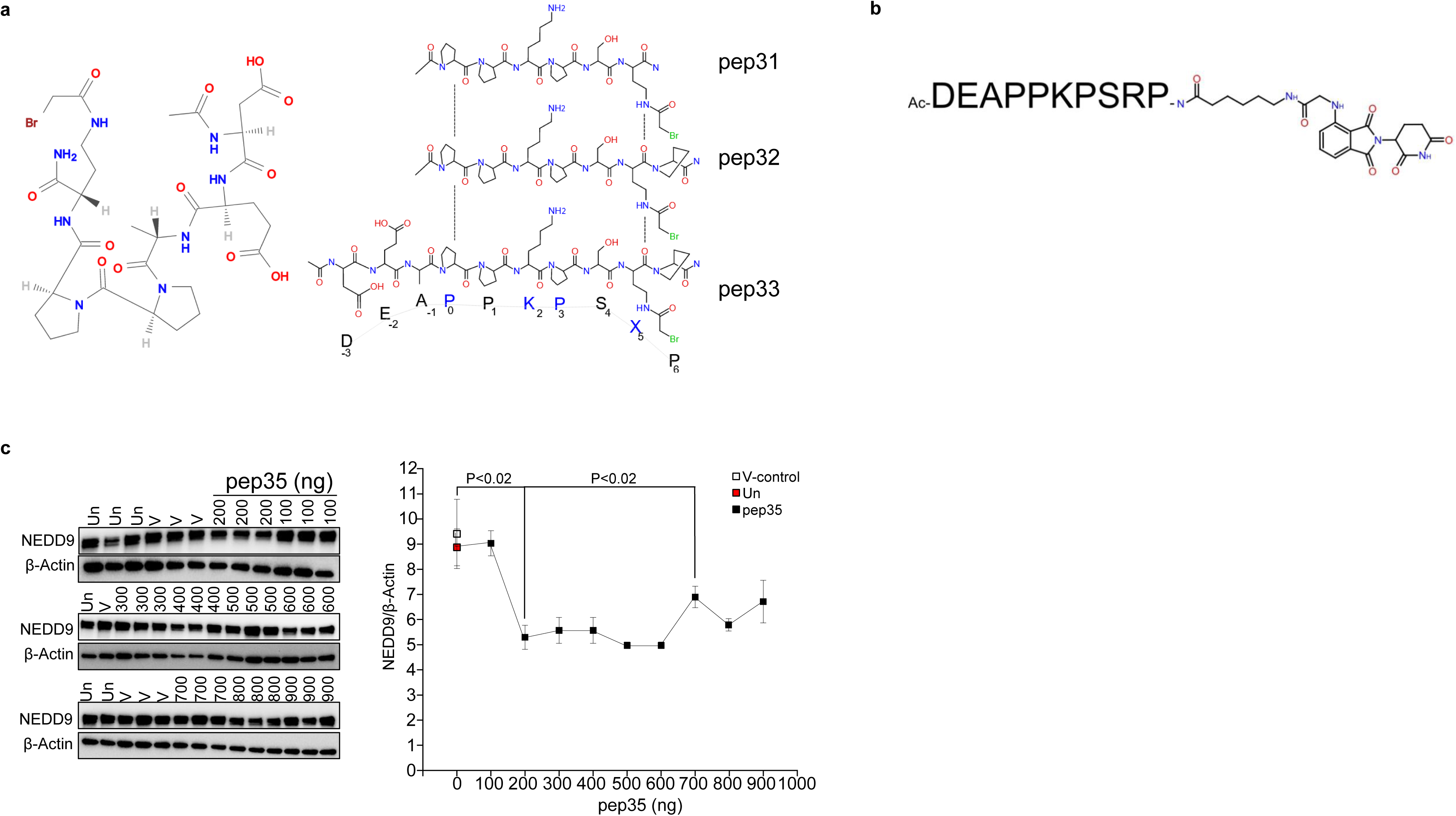
Degrader effect of NEDD9 by thalidomated NEDDtide. **a.** The chemical formula for the pep31, pep32, and pep32 series NEDDtides modified with a bromoacetamide residue. **b.** Structure of NEDDtide pep41 including thalidomide moiety. **c.** Cultured human pulmonary artery endothelial cells (HPAECs) were transfected with increasing doses of thalidomide-modified pep41 for 2 hr and NEDD9 expression was measured by immunoblot. The graph shows NEDD9/β-actin ratio for different concentrations of the peptide (*n*=3-5/condition).

**Fig. S8.**
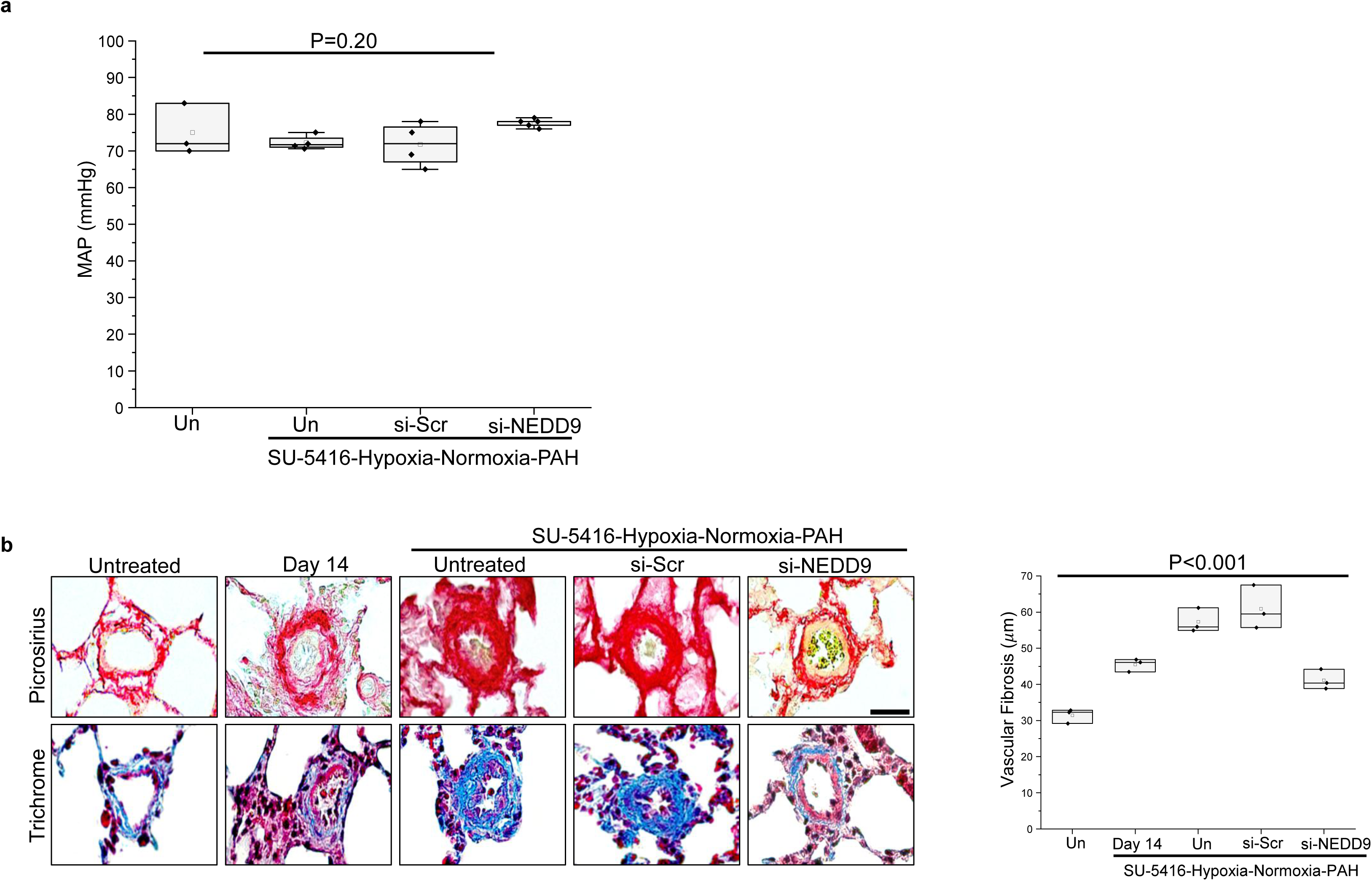
Molecular inhibition of NEDD9 with siRNA reverse vascular fibrosis in experimental PAH. In a disease-reversal treatment protocol, SU-5416-Hypoxia-Normoxia rats were administered siRNA against scrambled (negative) control (Scr) or NEDD9 beginning at protocol day 14, which is a timepoint associated with substantial pulmonary arterial remodeling. **a.** As expected, at the completion of the protocol there was no significant difference in mean arterial pressure across conditions measured by invasive cardiac catheterization (*n*=3-5/condition). Un, untreated. **b.** Paraffin-embedded lung sections were harvested from Sprague Dawley rats at various time points during the experimental protocol and analyzed using Masson trichome and Picrosirius red staining to measure global and fibrillar collagen, respectively, deposition in distal pulmonary arterioles measuring 25-50μm in diameter (*n*=3 rats/condition). Scale bar, 10 μm.

**Fig. S9.**
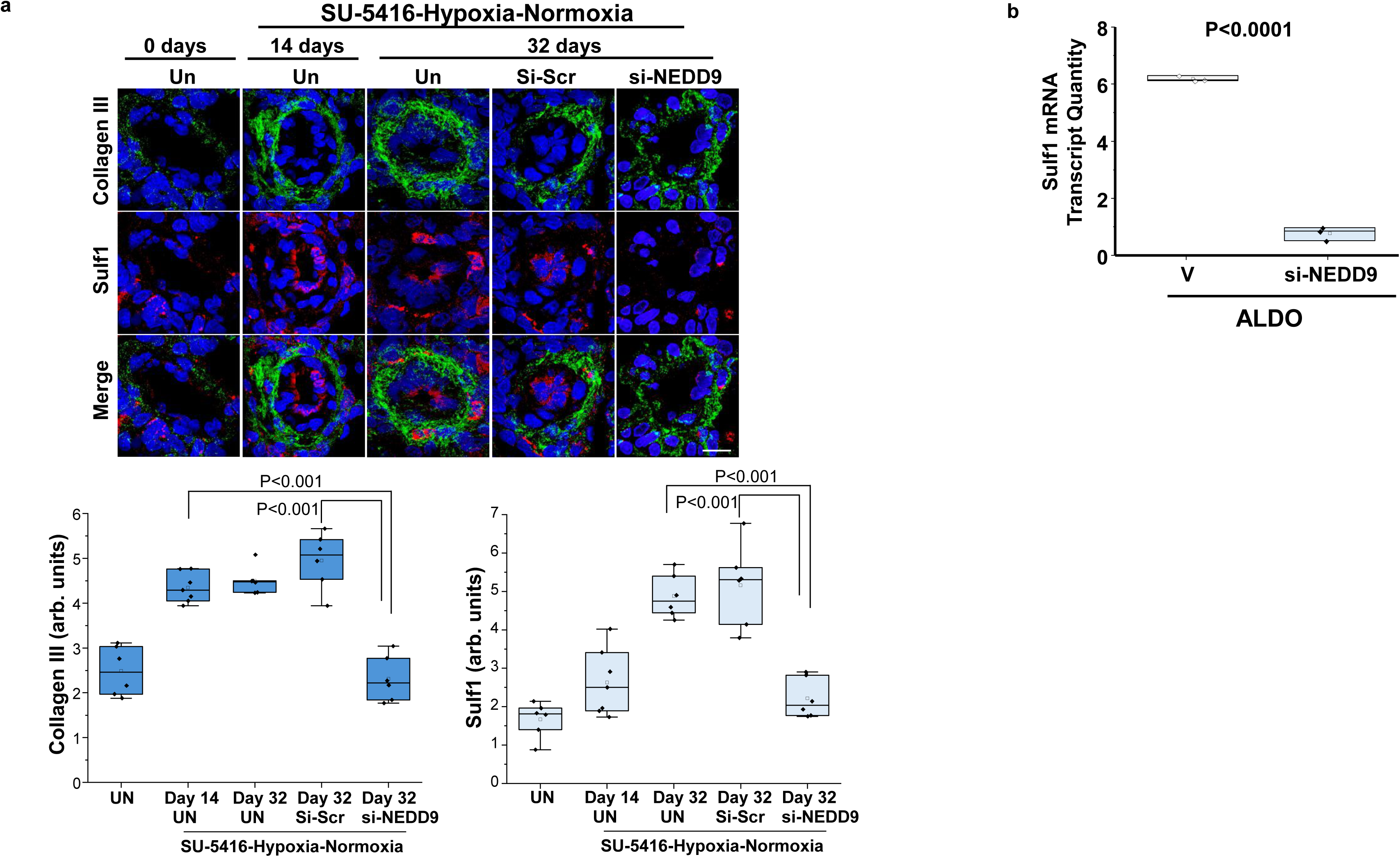
NEDD9 decreases Sulf1 in pulmonary arterial endothelial cells *in vitro* and remodeled pulmonary arterioles in PAH *in vivo*. **a.** Sprague Dawley rats were untreated (Un) or administered SU-5416-Hypoxia-Normoxia to induce PAH and treated with si-NEDD9 or si-scrambled (negative) control (Scr) per the disease reversal protocol illustrated in Fig. 3a. Paraffin-embedded lung sections were harvested and analyzed using anti-Sulf1 and anti-collagen III immunofluorescence (*n*=5-6 rats/condition). Blue, Dapi. Scale bar, 10 μm. **b.** Human pulmonary artery endothelial cells were treated with vehicle (V) control or transfected with siRNA against NEDD9 (si-NEDD9) for 24 hr and Sulf1 mRNA transcript quantity was analyzed by RT-qPCR. These data are extracted from findings reported originally in *reference* 8.

**Fig. 10.**
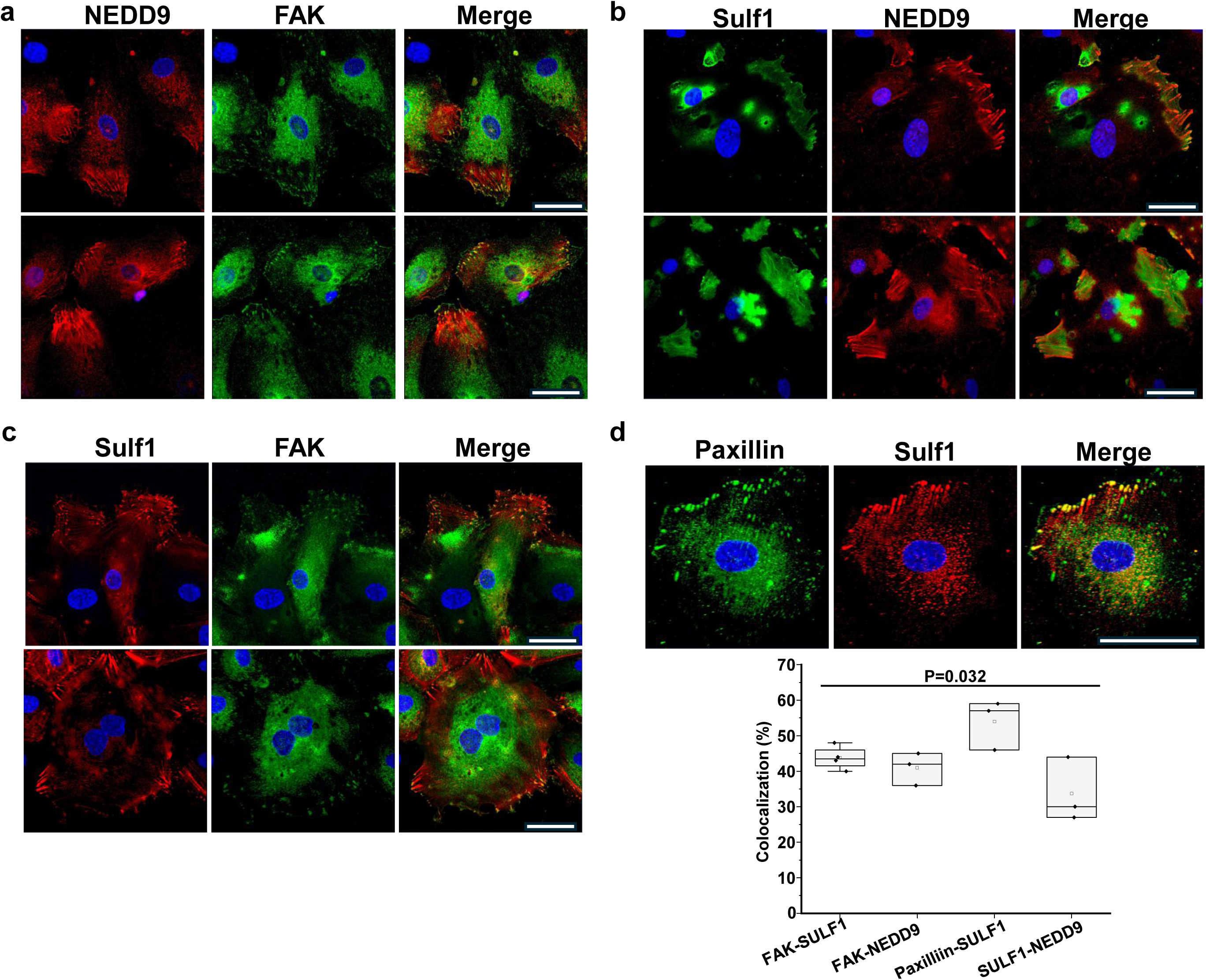
Co-localization of NEDD9, Sulf1, Paxillin, and FAK in cultured human pulmonary artery endothelial cells. Human pulmonary artery endothelial cells were analyzed using double immunofluorescence after probing with **a** anti-NEDD9 and anti-FAK, **b** anti-NEDD9 and anti-Sulf1, **c** anti-Sulf1 and anti-FAK, and **d** anti-Paxillin and anti Sulf1 antibodies (*n*=3-4/condition for all analysis). Blue, Dapi. FAK, focal adhesion kinase; Sulf1, sulfatase-1. Scale bar, 40 µm.

**Fig. S11.**
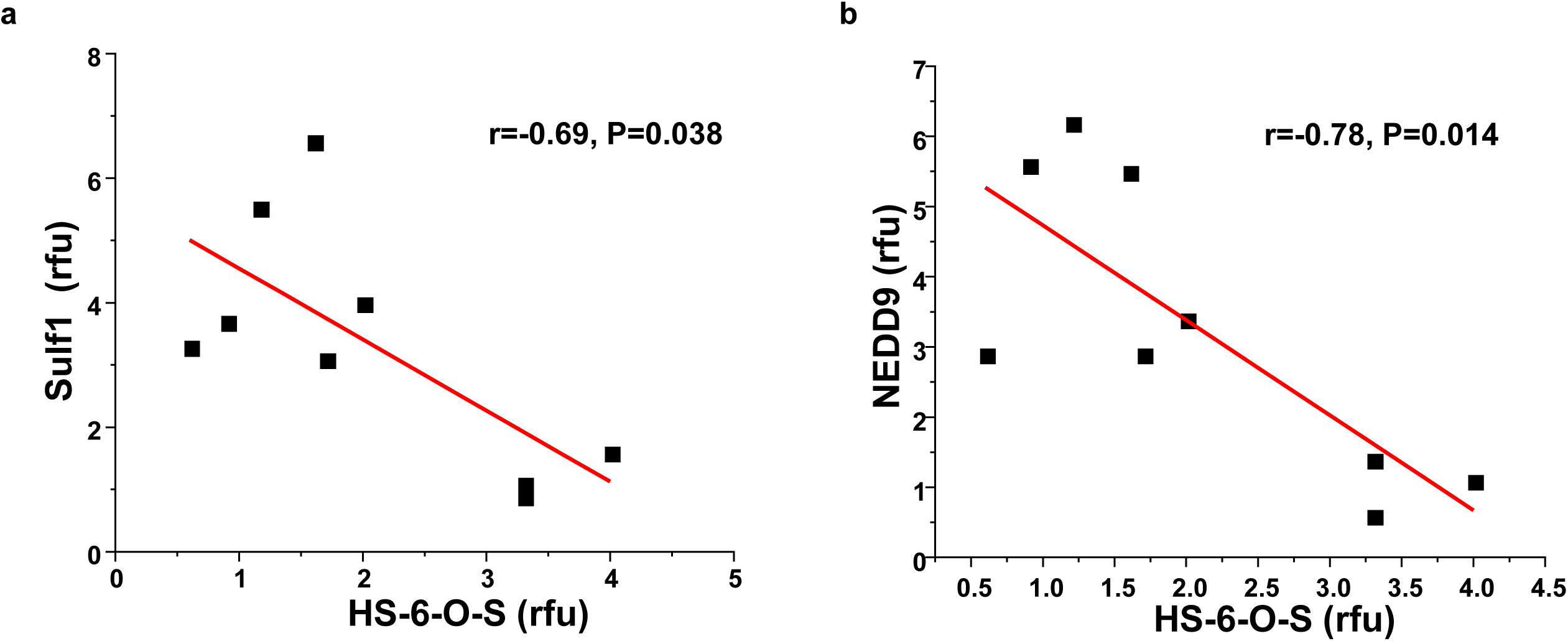
The relationship between pulmonary vascular heparan sulfate, NEDD9, and Sulf1 expression in PAH in vivo. **a.** Pearson correlation analysis comparing pulmonary vascular expression of 6-OS-Heparan sulfate (HS) with NEDD9 or **b.** Sulf1 in control, monocrotaline-PAH, and SU-5416-Hypoxia-Normoxia-PAH *in situ* (*n*=3 rats/condition). rfu, relative fluorescence unit. Comparisons were made using NEDD9 and Sulf1 data that were not measured together from within the same sample.

**Fig. S12.**
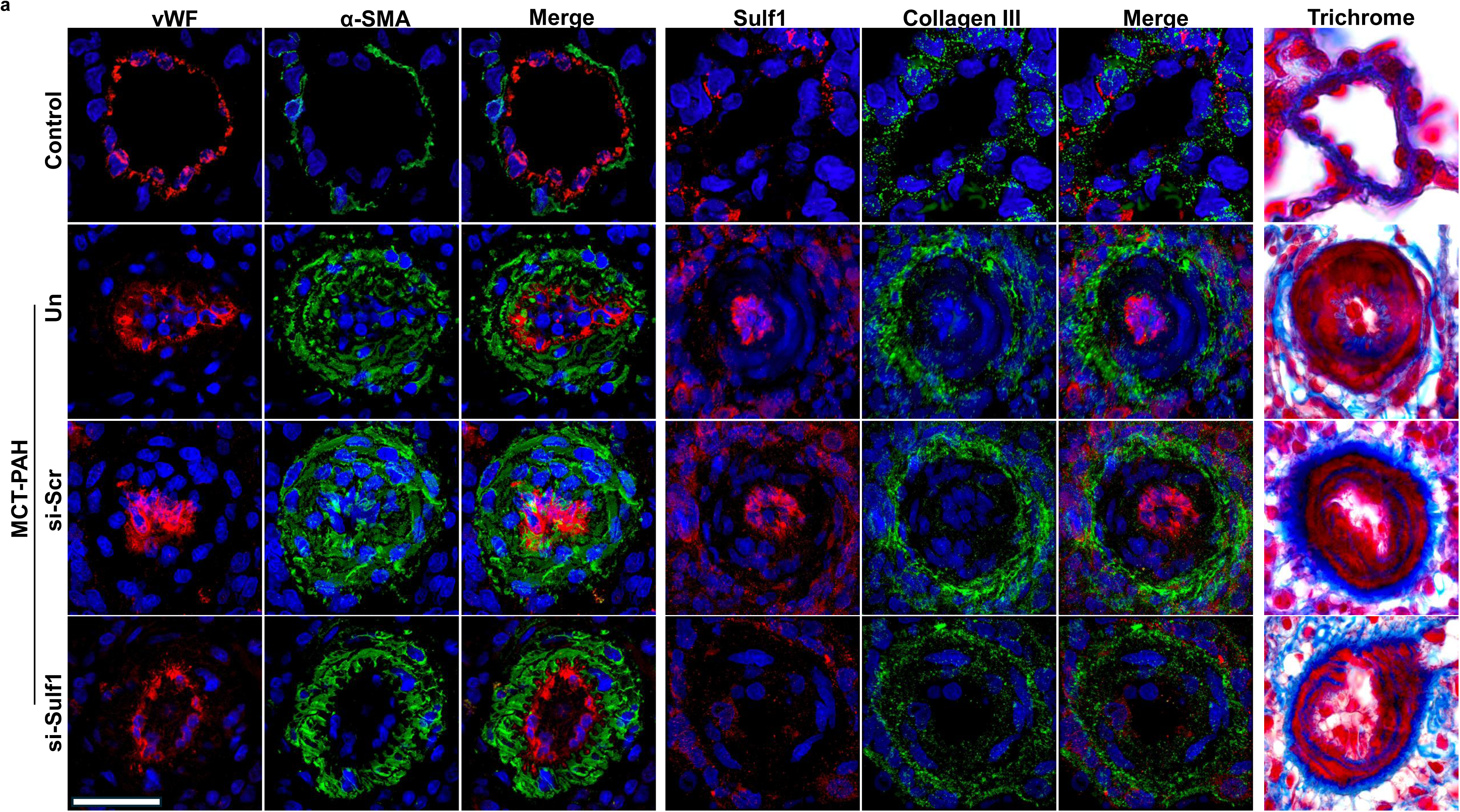
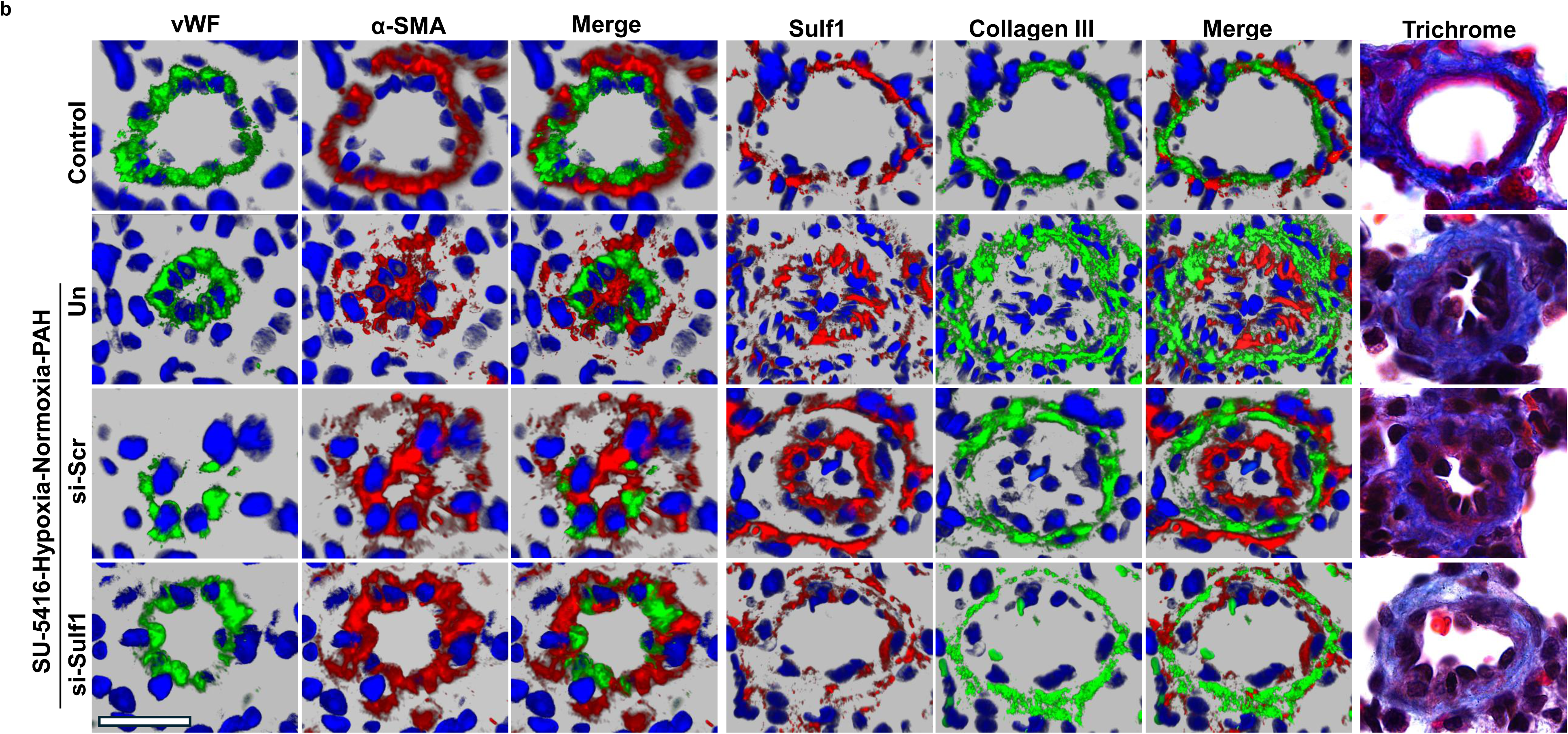
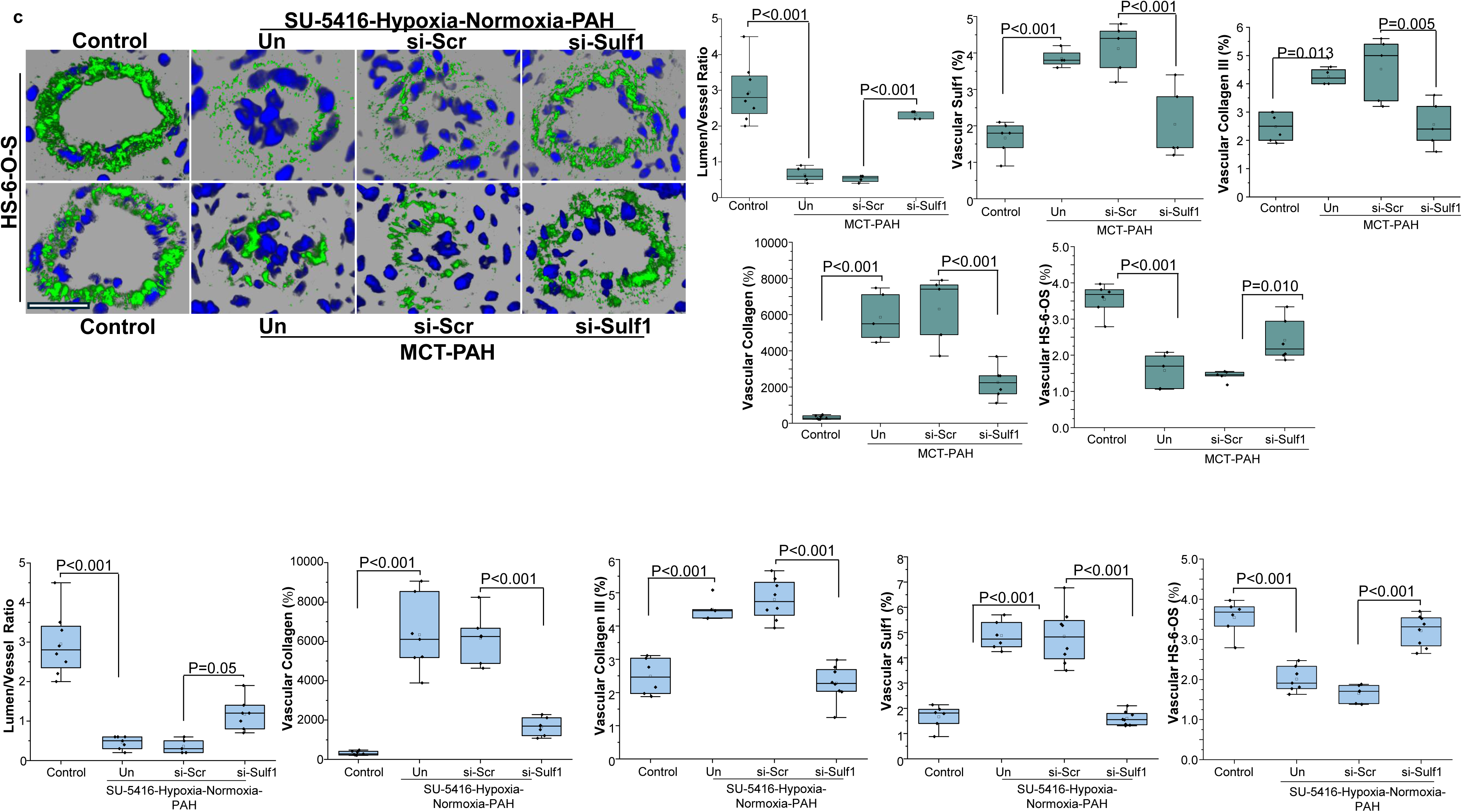
The effect of Sulf1 inhibition with siRNA on pulmonary vascular remodeling in two experimental models of PAH *in vivo*. In a disease-prevention treatment protocol, (**a**) monocrotaline (MCT)-PAH or (**b**) SU-5416-Hypoxia-Normoxia rats were administered siRNA against scrambled (negative) control (Scr) or sulfatase-1 (Sulf1) beginning at protocol day 1 and day 7, respectively. Paraffin-embedded lung sections were harvested from the rats at the completion of the protocol and stained with an antibody against von Willebrand Factor (vWF), alpha-smooth muscle actin (SMA), Sulf1, and collagen III or with with Masson trichrome, or (**c**) heparan sulfate-6-O-S (Blue, DAPI). The protein expression and fibrosis profile of distal pulmonary arterioles measuring 25-50μm in diameter was analyzed (*n=*4-8 rats/condition). Un, untreated. Scale bar, 20 μm.

**Fig. S13.**
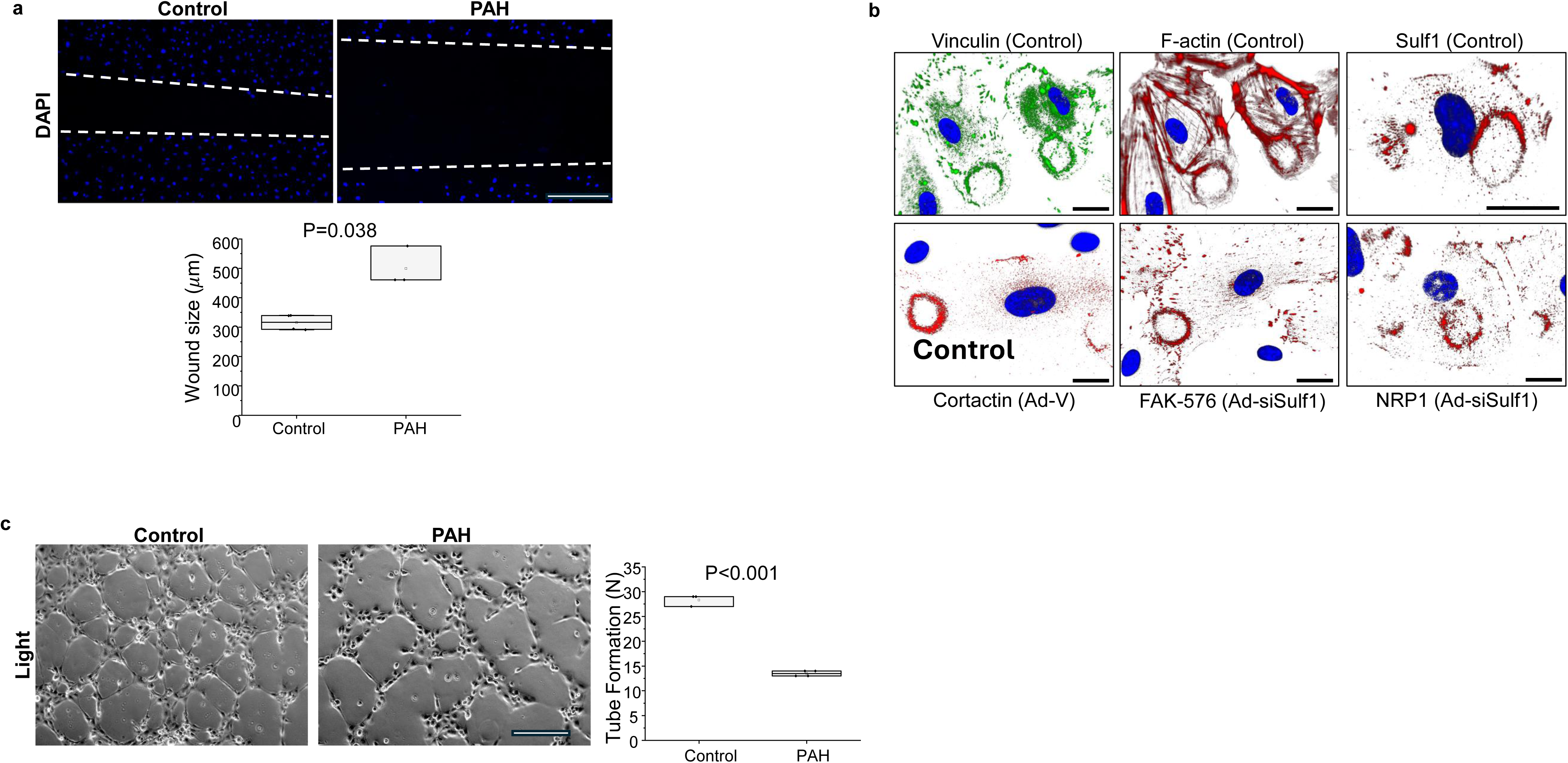
The pulmonary artery endothelial migration and podosome profile in pulmonary arterial hypertension (PAH). **a.** Compared to human pulmonary artery endothelial cells (HPAECs) from normal control donors, cell migration is decreased in PAH-HPAECs assessed by wound healing assay (*n*=3-4/condition). Scale bar, 400 μm. **b.** Cultured HPAECs were untreated or transfected with adenovirus (Ad) carrying vehicle (V) control or siRNA-against Sulf1 (si-Sulf1) and probed with antibodies against vinculin, F-actin, Sulf1, contracting, FAK-Y576, and NRP1. Scale bar, 20 μm. FAK, focal adhesion kinase; NRP1, neuropilin-1. (*n*=3-4/condition) **c.** Compared to HPAECs from normal control donors, endothelial tube formation is decreased in PAH (*n*=3-4/condition). Scale bar, 300 μm.

## Supplementary Tables

**Table S1.**
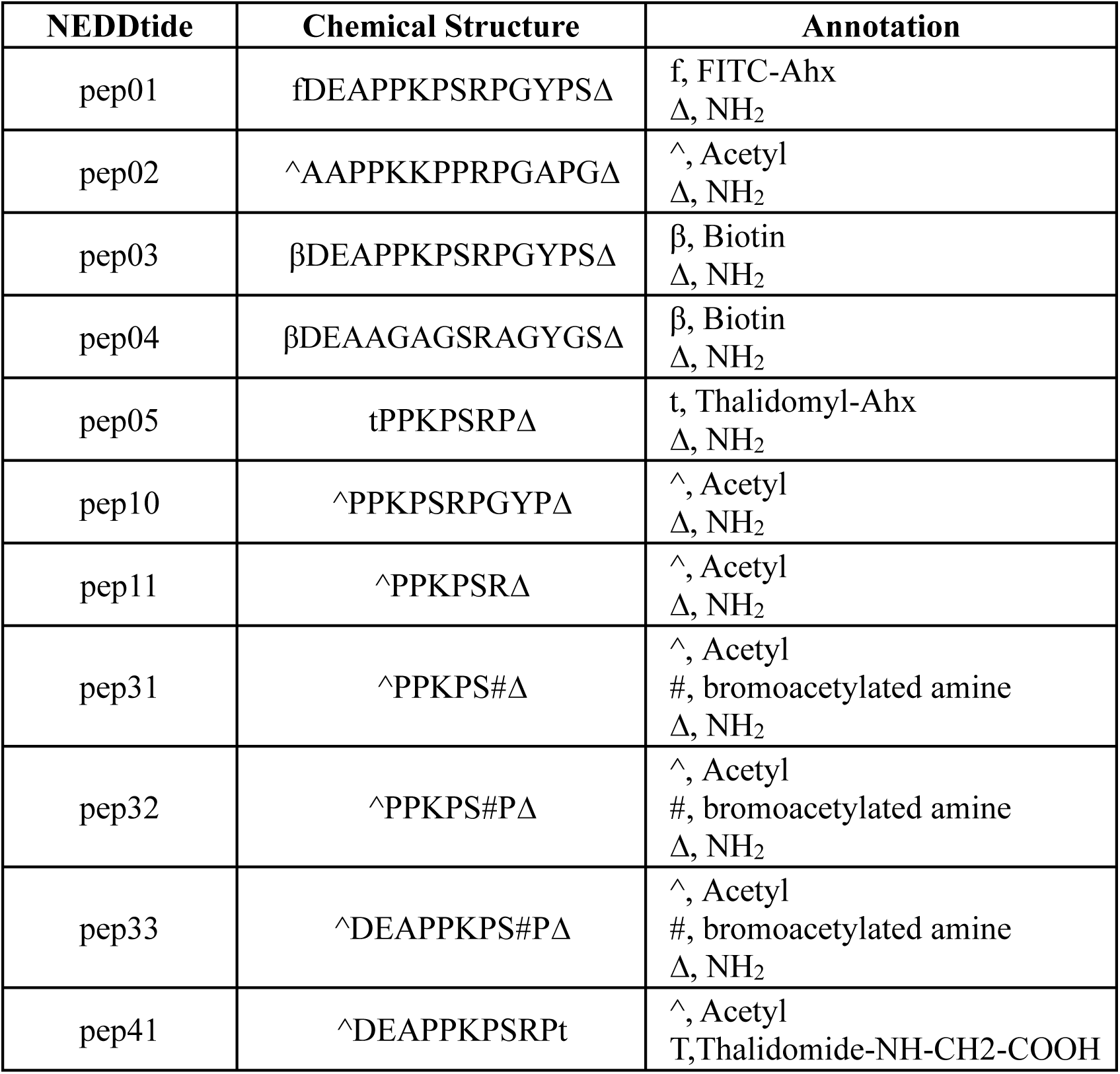
Catalogue of NEDDtides. FITC, Fluorescein isothiocyanate.

**Table S2.**
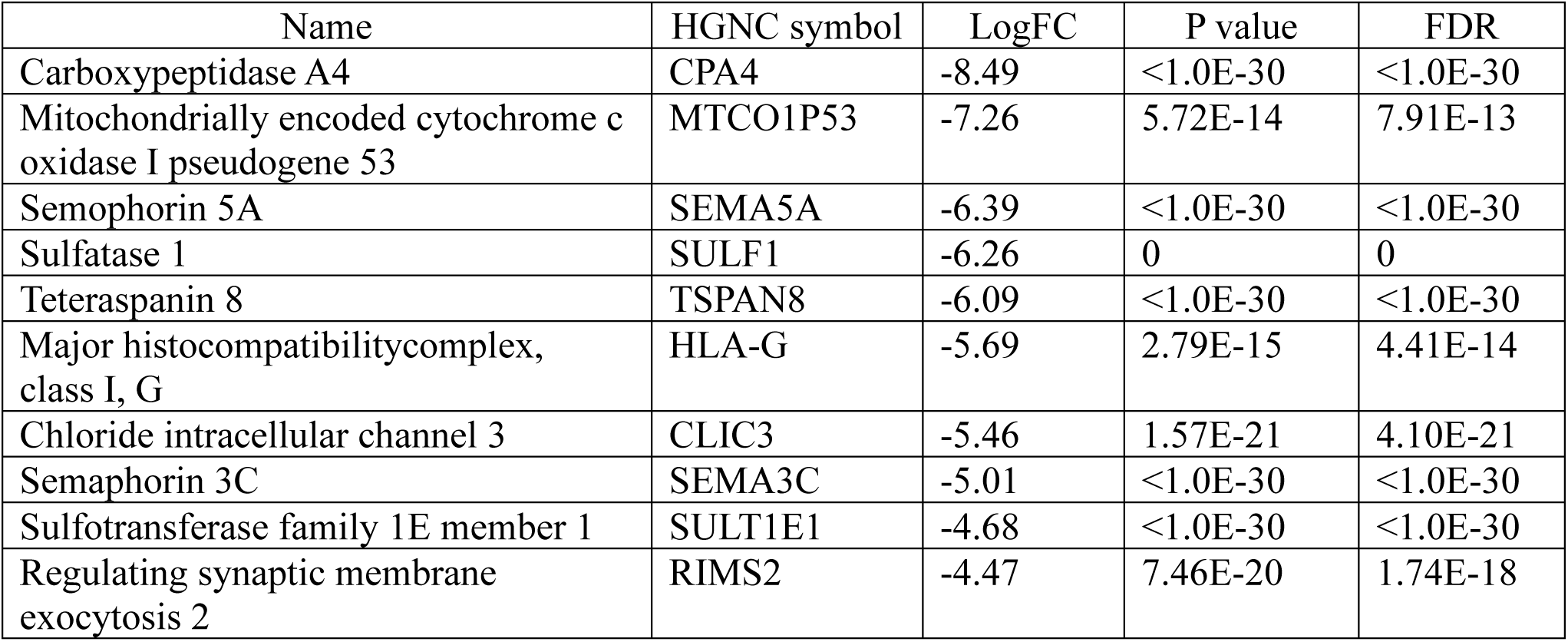
Top down-regulated genes regulated by NEDD9. Human pulmonary artery endothelial cells were untransfected or transfected with siRNA against NEDD9 and treated with the pro-oxidant stress pulmonary arterial hypertension hormone aldosterone and bulk RNA-Seq was performed. Results are derived from data reported originally in *ref.* 8.

| Name | HGNC symbol | LogFC | P value | FDR |
| --- | --- | --- | --- | --- |
| Carboxypeptidase A4 | CPA4 | -8.49 | <1.0E-30 | <1.0E-30 |
| Mitochondrially encoded cytochrome c oxidase I pseudogene 53 | MTCO1P53 | -7.26 | 5.72E-14 | 7.91E-13 |
| Semaphorin 5A | SEMA5A | -6.39 | <1.0E-30 | <1.0E-30 |
| Sulfatase 1 | SULF1 | -6.26 | 0 | 0 |
| Tetraspanin 8 | TSPAN8 | -6.09 | <1.0E-30 | <1.0E-30 |
| Major histocompatibility complex, class I, G | HLA-G | -5.69 | 2.79E-15 | 4.41E-14 |
| Chloride intracellular channel 3 | CLIC3 | -5.46 | 1.57E-21 | 4.10E-21 |
| Semaphorin 3C | SEMA3C | -5.01 | <1.0E-30 | <1.0E-30 |
| Sulfotransferase family 1E member 1 | SULT1E1 | -4.68 | <1.0E-30 | <1.0E-30 |
| Regulating synaptic membrane exocytosis 2 | RIMS2 | -4.47 | 7.46E-20 | 1.74E-18 |

**Table S3.**
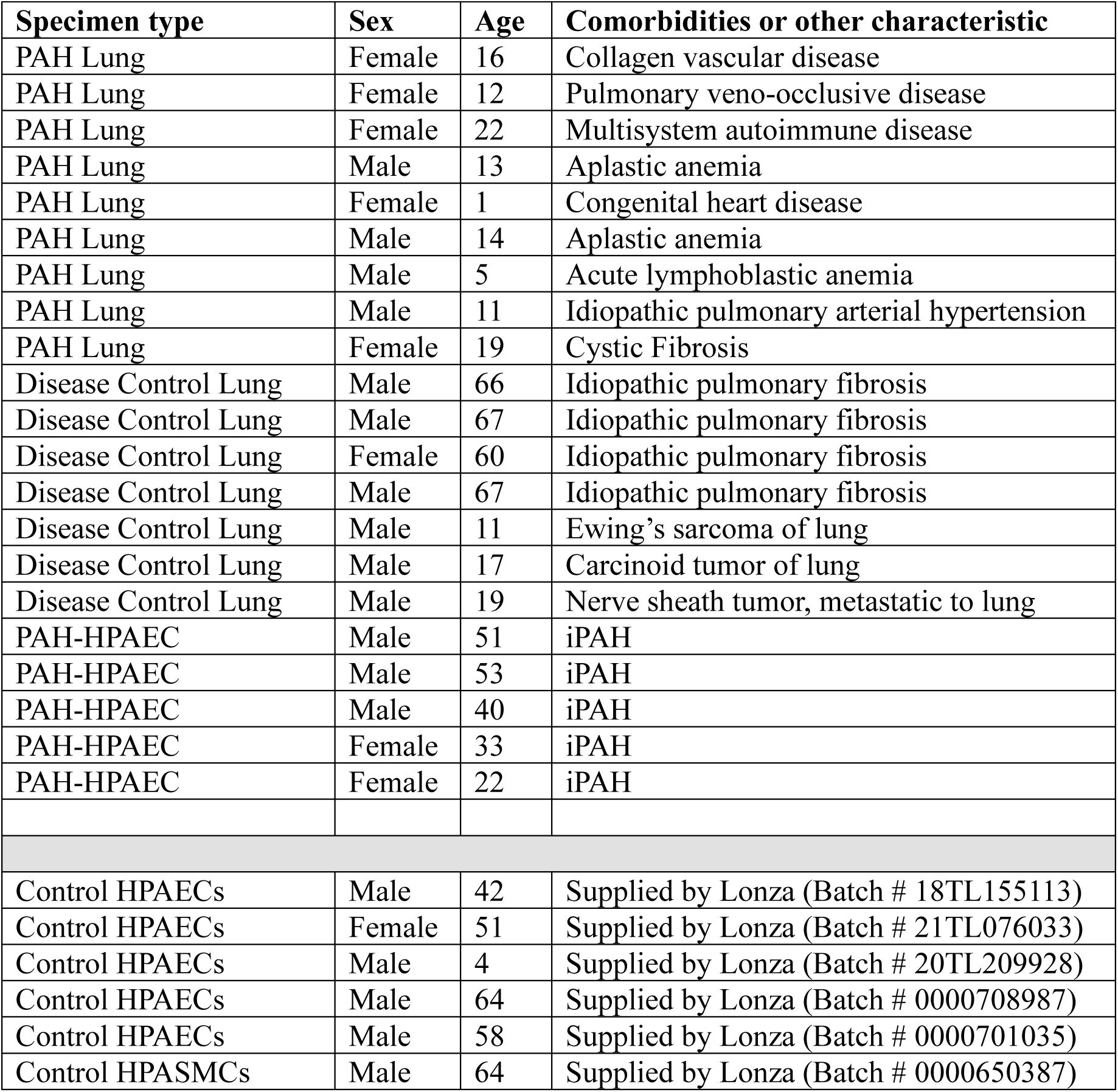
Inventory of biospecimens used in this study. iPAH, idiopathic pulmonary arterial hypertension; PAH-HPAEC, pulmonary arterial hypertension-human pulmonary artery endothelial cells; HPASMCs, human pulmonary artery smooth muscel cells.

**Table S4.**
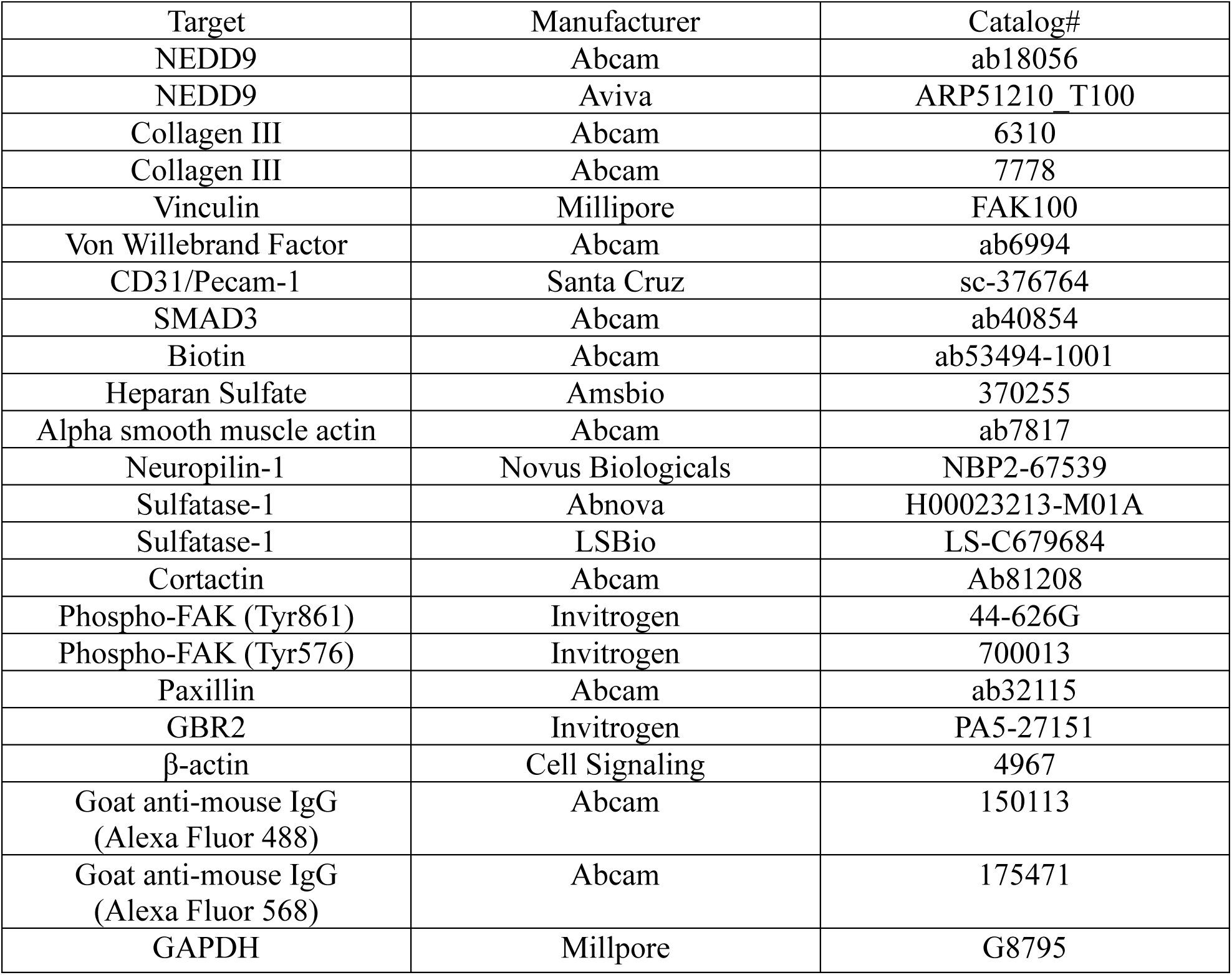
Inventory of antibodies and other commercially purchased reagents used in this study.

**Table S5.** Crystallographic data and refinement statistics for co-crystallization of NEDD9-SH3 domain with fragment 2.

|  | NEDD9/Apo | NEDD9/Apo(higher res) | BCAR1/FAKp10 | NEDD9(C18D)/Apo | NEDD9-Cpd | NEDD9/FAK1p33 |
| --- | --- | --- | --- | --- | --- | --- |
| Wavelength | 0.9792 | 0.9792 | 0.9792 | 0.9792 |  | 0.9202 |
| **Resolution range** | **43.55 - 2.6 (2.693 - 2.6)** | **44.8 - 2.28 (2.362 - 2.28)** | **33.82 - 1.2 (1.243 - 1.2)** | **31.99 - 1.77 (1.833 - 1.77)** | **46.1 - 2.204 (2.26 - 2.2)** | **54.15 - 1.91 (1.95 - 1.91)** |
| Space group | P 1 21 1 | P 61 2 2 | C 2 2 21 | P 1 21 1 | C 1 2 1 | P 1 21 1 |
| Unit cell | 52.87 48.83 98 90 100.62 90 | 51.73 51.73 429.83 90 90 120 | 38.73 69.36 57.43 90 90 90 | 51.83 48.61 84.95 90 93.02 90 | 92.21 53.15 167.27 90 90.07 90 | 60.03 38.2 124.08 90 90.3 90 |
| Total reflections | 50472 (4867) | 254500 (16916) | 144555 (8584) | 136582 (14032) | 76809 (5008) | 151972 (9357) |
| Unique reflections | 15066 (1519) | 16956 (1674) | 24202 (2154) | 40367 (3987) | 40680 (2683) | 44125 (2666) |
| Multiplicity | 3.4 (3.2) | 15.0 (10.1) | 6.0 (4.0) | 3.4 (3.5) | 1.0 (1.0) | 3.4 (3.5) |
| Completeness (%) | 97.39 (98.18) | 99.94 (99.76) | 98.46 (89.17) | 97.22 (96.95) | 98.47 (93.32) | 99.32 (99.21) |
| Mean I/sigma(I) | 9.12 (2.02) | 7.48 (0.71) | 13.91 (0.94) | 7.82 (0.81) | 5.28 (0.86) | 5.42 (0.69) |
| Wilson B-factor | 46.27 | 51.23 | 16.25 | 27.92 | 44.30 | 29.09 |
| R-merge | 0.1043 (0.5267) | 0.2233 (2.423) | 0.07932 (1.331) | 0.2025 (1.844) | 0.08278 (1.076) | 0.1547 (2.08) |
| R-meas | 0.1247 (0.6328) | 0.2311 (2.551) | 0.08753 (1.537) | 0.2438 (2.177) | 0.1123 (1.454) | 0.1836 (2.458) |
| R-pim | 0.06751 (0.3458) | 0.05836 (0.7778) | 0.03625 (0.7467) | 0.1336 (1.145) | 0.07527 (0.9691) | 0.09795 (1.299) |
| CC1/2 | 0.992 (0.925) | 0.996 (0.519) | 0.998 (0.367) | 0.972 (0.257) | 0.994 (0.704) | 0.995 (0.367) |
| CC* | 0.998 (0.98) | 0.999 (0.827) | 0.999 (0.733) | 0.993 (0.639) | 0.999 (0.909) | 0.999 (0.733) |
| Reflections used in refinement | 15023 (1511) | 16948 (1671) | 24192 (2150) | 40313 (3978) | 40680 (2683) | 44017 (2652) |
| Reflections used for R-free | 761 (95) | 860 (78) | 1250 (112) | 1893 (224) | 1990 (131) | 2226 (103) |
| R-work | 0.2356 (0.3494) | 0.2648 (0.3989) | 0.1903 (0.3999) | 0.2359 (0.4243) | 0.2395 (0.4245) | 0.2438 (0.4102) |
| **R-free** | **0.2905 (0.4082)** | **0.2965 (0.4207)** | **0.2104 (0.3573)** | **0.2827 (0.4872)** | **0.2920 (0.4867)** | **0.2754 (0.4401)** |
| CC(work) | 0.940 (0.792) | 0.942 (0.640) | 0.959 (0.657) | 0.951 (0.590) |  |  |
| CC(free) | 0.930 (0.554) | 0.917 (0.461) | 0.950 (0.643) | 0.917 (0.410) |  |  |
| Number of non-hydrogen atoms | 3454 | 1939 | 668 | 3559 | 4355 | 4019 |
| macromolecules | 3383 | 1872 | 589 | 3290 | 4252 | 3784 |
| ligands | 0 | 0 | 0 | 0 | 78 | 1 |
| solvent | 71 | 67 | 79 | 269 | 25 | 234 |
| Protein residues | 449 | 240 | 75 | 420 | 548 | 487 |
| Nucleic acid bases | 0 | 0 |  |  |  |  |
| RMS(bonds) | 0.004 | 0.005 | 0.010 | 0.002 | 0.007 | 0.013 |
| RMS(angles) | 1.27 | 1.21 | 1.30 | 0.56 | 0.97 | 1.44 |
| Ramachandran favored (%) | 99.54 | 99.14 | 94.29 | 99.75 | 93.58 | 99.78 |
| Ramachandran allowed (%) | 0.46 | 0.86 | 5.71 | 0.25 | 5.66 | 0.22 |
| Ramachandran outliers (%) | 0.00 | 0.00 | 0.00 | 0.00 | 0.75 | 0.00 |
| Rotamer outliers (%) | 1.80 | 1.53 | 0.00 | 1.75 | 1.84 | 1.01 |
| Clashscore | 6.02 | 9.31 | 8.45 | 6.38 | 7.52 | 5.19 |
| Average B-factor | 52.16 | 61.15 | 21.46 | 35.12 | 57.26 | 39.15 |
| macromolecules | 52.25 | 61.32 | 20.22 | 34.72 | 56.77 | 38.76 |
| ligands | 0 | 0 |  |  | 83.37 | 53.03 |
| solvent | 48.28 | 56.37 | 30.70 | 40.07 | 59.11 | 41.06 |
| Number of TLS groups | 22 | 1 |  | 21 |  |  |

Values in parentheses are for highest resolution shell RMS = root mean square

## References

1. Mayer, B.J. The discovery of modular binding domains: building blocks of cell signaling. Nat Rev Mol Cell Biol 16, 855–858 (2015).

2. Moarefi, I., et al. Activation of the Src-family tyrosine kinase Hck by SH3 domain displacement. Nature 385, 650–3 (1997).

3. Law, S.F., et al. Human enhancer of filamentation 1, a novel p130cas-like docking protein, associates with focal adhesion kinase and induces pseudohyphal growth in Saccharomyces cerevisiae. Mol Cell Biol 16, 3327–37 (1996).

4. O’Neill, G.M., Golemis, E.A. Proteolysis of the docking protein HEF1 and implications for focal adhesion dynamics. Mol Cell Biol 21, 5094–108 (2001).

5. Faure, A.J., et al. Mapping the energetic and allosteric landscapes of protein binding domains. Nature 604, 1775–83 (2022).

6. Lu, H., et al. Recent advances in the development of protein–protein interactions modulators: mechanisms and clinical trials. Sig Transduct Target Ther 5, 213 (2020).

7. Bradbury, P., Bach, C.T., Paul, A., O’Neill, G.M. Src kinase determines the dynamic exchange of the docking protein NEDD9 (neural precursor cell expressed developmentally down-regulated gene 9) at focal adhesions. J Biol Chem 289, 24792–800 (2014).

8. Samokhin, A.O., et al. NEDD9 targets COL3A1 to promote endothelial fibrosis and pulmonary arterial hypertension. Sci Transl Med 445, eaap7294 (2018).

9. Wisniewska, M., et al. The 1.1 A resolution crystal structure of the p130cas SH3 domain and ramifications for ligand selectivity. J Mol Biol 347, 1005–14 (2005).

10. Maron, B.A. Revised definition of pulmonary hypertension and approach to management: A clinical primer. J Am Heart Assoc 12, e029024 (2023).

11. Niihori, M., et al. Mitochondria as a primary determinant of angiogenic modality in pulmonary arterial hypertension. J Exp Med 221, e20231568 (2024).

12. Grobs, Y., et al. ATP citrate lyase drives vascular remodeling in systemic and pulmonary vascular diseases through metabolic and epigenetic changes. Sci Transl Med 16, eado7824 (2024).

13. Wang, H.B., Dembo, M., Hanks, S.K., Wang, Y. Focal adhesion kinase is involved in mechanosensing during fibroblast migration. Proc Natl Acad Sci USA 98, 11295–300 (2001).

14. Paulin, R., et al. Targeting cell motility in pulmonary arterial hypertension. Eur Respir J 43, 531–44 (2014).

15. Oshima, K., et al. Loss of endothelial sulfatase-1 after experimental sepsis attenuates subsequent pulmonary inflammatory responses. Am J Physiol Lung Cell Mol Physiol 317, L667–L677 (2019).

16. Justo, T., Martiniuc, A., Dhoot, G.K. Modulation of cell signaling and sulfation in cardiovascular development and disease. Sci Rep 11, 22424 (2021).

17. Korf-Klingebiel, M., et al. Heparan sulfate-editing extracellular sulfatases enhance vegf bioavailability for ischemic heart repair. Circ Res 125,787–801 (2019).

18. Lu, J., Auduong, L., White, E.S., Yue, X. Up-regulation of heparan sulfate 6-O-sulfation in idiopathic pulmonary fibrosis. Am J Respir Cell Mol Biol 50, 106–114 (2014).

19. Masola V, Greco N, Gambaro G, Franchi M, Onisto M. Heparanase as active player in endothelial glycocalyx remodeling. Matrix Biol Plus 2021 Dec 25:13:100097.

20. Moon JJ, Matsumoto M, Patel S, Lee L, Guan JL, Li S. Role of cell surface heparan sulfate proteoglycans in endothelial cell migration and mechanotransduction. J Cell Physiol 2005 Apr;203(1):166–76.

21. Goncharova, E.A., Kudryashova, T.V., de Jesus Perez, V.A., Rafikova, O. UnWNTing the heart: targeting wnt signaling in pulmonary arterial hypertension. Circ Res 132, 1486–88 (2023).

22. Yuan, K, et al. Loss of endothelium-derived wnt5a is associated with reduced pericyte recruitment and small vessel loss in pulmonary arterial hypertension. Circulation 139, 1710–24 (2019).

23. Humbert, M., et al. Pathology and pathobiology of pulmonary hypertension: state of the art and research perspectives. Eur Respir J 53, 1801887 (2019).

24. Backus, K.M., et al. Proteome-wide covalent ligand discovery in native biological systems. Nature 534, 570–4 (2016).

25. Teyra J, et al. Comprehensive analysis of the human SH3 domain family reveals a wide variety of noncanonical specificities. Structure 2017;25:1598–610.

26. Sigrist, C.J.A., et al. New and continuing developments at PROSITE Nucl Acids Res 41, D344–D347 (2013).

27. Weisberg, E.L., et al. Inhibition of USP10 induces degradation of oncogenic FLT_3_. Nat Chem Biol 13, 1207–15 (2017).

28. Xie, P., et al. A fiducial-assisted strategy compatible with resolving small MFS transporter structures in multiple conformations using cryo-EM. Nat Commun 16, 7 (2025).

29. Marchant, C.L., Malmi-Kakkada, A.N., Espina, J.A., & Barriga, E.H. Cell clusters softening triggers collective cell migration in vivo. Nat Mat 21 1314–1323 (2022).

30. Niihori, M., et al. Mitochondria as a primary determinant of angiongenic modality in pulmonary arterial hypertension. J Exp Med 221, e20231568 (2024).

31. Maron, B.A., et al. Plasma aldosterone levels are elevated in patients with pulmonary arterial hypertension in the absence of left ventricular heart failure: a pilot study. Eur J Heart Fail 15, 277–83 (2013).

32. Tran, T.H., Shi, X., Zaia, J., Ai, X. Heparan sulfate 6-O-endosulfatases (Sulfs) coordinate the Wnt signaling pathways to regulate myoblast fusion during skeletal muscle regeneration. J Biol Chem 287,32651–64 (2012).

33. Abe, K., et al. Formation of plexiform lesions in experimental severe pulmonary arterial hypertension. Circulation 121, 2747–54 (2010).

34. Tan, X., et al. Focal adhesion kinase: from biological functions to therapeutic strategies. Exp Hematol Oncol 12, 83 (2023).

35. Pan, Y.R., Chen, C.L., Chen, H.C. FAK is required for the assembly of podosome rosettes J Cell Biol 195, 113–29 (2011).

36. Seano, G., et al. Endothelial podosome rosettes regulate vascular branching in tumour angiogenesis. Nat Cell Biol 16, 931–41 (2014).

37. Lim, W.A., Richards, F.M., Fox, R.O. Structural determinants of peptide-binding orientation and of sequence specificity in SH3 domains. Nature 372, 375–379 (1994).

38. Tikhmyanova, N., Little, J.L., Golemis, E.A. CAS proteins in normal and pathological cell growth cycle. Cell Mol Life Sci 67, 1025**-**1048 (2009).

39. Tu, L., et al. A critical role for p130Cas in the progression of pulmonary hypertension in humans and rodents. Am J Crit Care Med 186, 666–76 (2012).

40. Santiago, A., Morano, K.A. Oxidation of two cysteines within yeast Hsp70 impairs proteostasis while directly triggering an Hsf1-dependent cytoprotective response. J Biol Chem 298, 102424 (2022).

41. Hwang, P.M., Bishop, R.E., Kay, L.E. The integral membrane enzyme PagP alternates between two dynamically distinct states. Proc Natl Acad Sci U S A. 101, 9618–23 (2004).

42. Muller, J.B., Geyer, P.E., Calaco, A.R., et al. The proteome landscape of the kingdoms of life. Nature 582, 592–96 (2020).

43. Singh, M., Cowell, L., Seo, S., O’Neill G., Golemis, E. Molecular basis for HEF1/NEDD9/Cas-L action as a multifunctional co-ordinator of invasion, apoptosis and cell cycle. Cell Biochem Biophys 48, 54–72 (2007).

44. Dieffenbach P.B., Haeger, C.M., Rehman, R., et al. A novel protective role for matrix metalloproteinase-8 in pulmonary vasculature. Am J Respir Crit Care Med 204, 1433–51 (2021).

45. Bradbury, P., Bach, C.T., Paul, A., O’Neill G.M. Src kinase determines the dynamic exchange of the docking protein NEDD9 (neural precursor cell expressed developmentally down-regulated gene 9) at focal adhesions. J Biol Chem 289, 24792–800 (2014).

46. Yang, Z., Chen, M., Ge, R., et al. Identification of a non-inhibitory aptameric ligand to CRL2^ZYG11B^ E3 ligase for targeted protein degradation. Nat Commun 16, 2494 (2025).

47. Omata, Y., Nakamura, S., Koyama, T., et al. Identification of Nedd9 a TGF-*#*-Smad2/3 target gene involved in RNAKL-induced osteoclastogenesis by comprehensive analysis. PLoS One 11, e-157992 (2016).

48. Izumchenko, E., Singh, M.K., Plotnikova, O.V., et al. NEDD9 promotes oncogenic signaling in mammary tumor development. Cancer Res 69, 7198–206 (2009).

49. Katayose, T., Iwata, S., Oyaizu, N., et al. The role of Cas-L/NEDD9 as a regulator of collagen-induced arthritis in a murine model. Biochem Biophys Res Commun 460, 1069–75 (2015).

50. Vattulainen-Collanus, S., Akinrinade, O., Li, M., et al. Loss of PPARγ in endothelial cells leads to impaired angiogenesis. J Cell Sci 129, 693–705 (2016).

51. Niihori, M., James, J., Varghese M.V., et al. Mitochondria as a primary determinant of angiogenic modality in pulmonary arterial hypertension. J Exp Med 222, e20231568 (2024).

52. Cober, N.D., VandenBroek, M.M., Ormiston, M.L., Stewart, D.J. Evolving concepts of endothelial pathobiology of pulmonary arterial hypertension. Hypertension 79, 1580–90 (2022).

53. Mutgan, A-C., Radic, N., Valzano, F., et al. A comprehensive map of proteoglycan expression and deposition in the pulmonary arterial wall in health and pulmonary hypertension. Am J Physiol Lung Cell Mol Physiol 327, L173–188 (2024).

54. Chang, Y-T., Chan, C.K., Eriksson, I., et al. Versican accumulates in vascular lesions in pulmonary arterial hypertension. Pulm Circ 6, 347–59 (2016).

55. Shimizu, M., Murakami, T., Suto, F., Fujisawa, H. Determination of cell adhesion sites of Neuropilin-1. J Cell Biol 148,1283–93 (2020).

56. Weiss, R.J., Spahn, R.N., Toledo, A.G., et al. ZNF263 is a transcriptional regulator of heparin and heparan sulfate biosynthesis. Proc Natl Acad Sci USA 117, 19311–17 (2020).

57. McGowan, S.E., Lankasara, T.I., McCoy, D.M., Zhu, L., Tivanski, A.V. Platelet-derived growth factor-α and neuropilin-1 mediate lung fibroblast response to rigid collagen fibers. Am J Respir Cell Mol Biol 62, 454–465 (2020).

58. Seano, G., Chiaverina, G., Gagliardi, P.A., et al. Endothelial podosome rosettes regulate vascular branching in tumour angiogenesis. Nat Cell Biol 16, 931–41 (2014).

59. Sulzmaier, F.J., Jean, C., Schlaepfer, D.D. FAK in cancer: mechanistic findings and clinical applications. Nat Rev Cancer 14, 598–610 (2014).

60. Wertheim, B.M., Wang, R.S., Guillermier, C., et al. Proline and glucose metabolic reprogramming supports vascular endothelial and medial biomass in pulmonary arterial hypertension. JCI Insight 8, e163932 (2023).

61. Steinhauser, M.L, Maron, B.A. Viewing pulmonary arterial hypertension pathogenesis and opportunities for disease-modifying therapy through the lens of biomass. JACC Basic Transl Sci 9, 1252–63 (2024).

## References

1. Backus, K.M., et al. Proteome-wide covalent ligand discovery in native biological systems. Nature 534:570–574 (2016).

2. Samokhin, A.O., et al. NEDD9 targets COL3A1 to promote endothelial fibrosis and pulmonary arterial hypertension. Sci Transl Med 445, eaap7294 (2018).

3. Kabsch, W. XDS. Acta Cryst D66:125–132 (2010).

4. Winter, G., et al. DIALS as a tookit. Protein Sci 31:232–250 (2021).

5. McCoy, A.J., et al. Phaser crystallographic software. J Appl Crystal 40:658–74 (2007).

6. Adams, P.D., et al. PHENIX: a comprehensive Python-based system for macromolecular structure solution. Acta Crys D66:213–21 (2010).

7. Afonine, P.V., et al. Real-space refinement in PHENIX for cryo-EM and crystallography. Acta Crystallogr D Struct Biol 74:531–544 (2018).

8. Emsley, P., Cowtan, K. Coot: model-building tools for molecular graphics. Acta Crys D60:2126–32 (2004).

9. Jensen, H., Ostergaard, J. Flow induced dispersion analysis quantifies noncovalent interactions in nanoliter samples. J Am Chem Soc 132:4070–1 (2010).

10. Jiang, J., et al. Discovery of covalent MKK4/7 dual inhibitor. Cell Chem Biol 27:1553–60 (2020).

11. Mori et al. Improved sensitivity of HSQC spectra of exchanging protons at short interscan delays using a new fast HSQC (FHSQC) detection scheme that avoids water saturation. J Magn Reson B. 108:94–8 (1995)

12. Jumper, J., et al. Highly accurate structure prediction with AlphaFold. Nature 596:583–89 (2021).

13. Maron, B.A., et al. Aldosterone inactivates the endothelin-B receptor via a cysteinyl thiol redox switch to decrease pulmonary endothelial nitric oxide levels and modulate pulmonary arterial hypertension. Circulation 126:963–74 (2012).

